# Presence and Transmission of Mitochondrial Heteroplasmic Mutations in Human Populations of European and African Ancestry

**DOI:** 10.1101/2020.10.13.337071

**Authors:** Chunyu Liu, Jessica L. Fetterman, Yong Qian, Xianbang Sun, Kaiyu Yan, Thomas Blackwell, Achilleas Pitsillides, Brian E. Cade, Heming Wang, Laura M. Raffield, Leslie A. Lange, Pramod Anugu, Goncalo Abecasis, L Adrienne Cupples, Susan Redline, Adolfo Correa, Ramachandran S Vasan, James Wilson, Jun Ding, Daniel Levy, NHLBI Trans-Omics for Precision Medicine (TOPMed) Consortium

## Abstract

We investigated the concordance of mitochondrial DNA heteroplasmic mutations (heteroplasmies) in different types of maternal pairs (n=6,745 pairs) of European (EA, n=4,718 pairs) and African (AA, n=2,027 pairs) Americans with whole genome sequences (WGSs). The average concordance rate of heteroplasmies was highest between mother-offspring pairs, followed by sibling-sibling pairs and more distantly related maternal pairs in both EA and AA participants. The allele fractions of concordant heteroplasmies exhibited high correlation (R^2^=0.8) between paired individuals. Compared to concordant heteroplasmies, discordant ones were more likely to locate in coding regions, be nonsynonymous or nonsynonymous-deleterious (p<0.001). The average number of heteroplasmies per individual (i.e. heteroplasmic burden) was at a similar level until older age (70-80 years old) and increased significantly thereafter (p<0.01). The burden of deleterious heteroplasmies (combined annotation-dependent depletion score≥15), however, was significantly correlated with advancing age (20-44, 45-64, ≥65 years, p-trend=0.01). A genome-wide association analysis of the heteroplasmic burden identified many significant (P<5e-8) common variants (minor allele frequency>0.05) at 11p11.12. Many of the top SNPs act as strong long-range cis regulators of protein tyrosine phosphatase receptor type J. This study provides further evidence that mtDNA heteroplasmies may be inherited or somatic. Somatic heteroplasmic variants increase with advancing age and are more likely to have an adverse impact on mitochondrial function. Further studies are warranted for functional characterization of the deleterious heteroplasmies occurring with advancing age and the association of the 11p11.12 region of the nuclear genome with mtDNA heteroplasmy.

## INTRODUCTION

Mitochondria are key organelles for energy metabolism, and they play a critical role in a variety of human diseases.^1^ The maternally inherited mitochondrial genome (mtDNA) is present in hundreds or thousands of copies in a cell, depending upon the cell type and its energetic needs. The mtDNA is a 16.6 kb double-stranded DNA that encodes 13 key subunits of the energy-producing oxidative phosphorylation (OXPHOS) pathway, as well as 22 transfer RNAs (tRNAs) and two ribosomal RNAs (rRNAs) for mitochondrial translation.^2^ A high mutation rate and maternal inheritance of the mtDNA has given rise to multiple mtDNA haplogroups that reflect the sequential accumulation of mtDNA polymorphisms (i.e. homoplasmic variants), revealing ancestry and patterns of prehistoric migration.^3^

Due to the presence of many mtDNA copies within a cell, heteroplasmic mutations (or heteroplasmies) may arise. Heteroplasmy is a phenomenon characterized by two or more mtDNA alleles co-existing at the same locus in different copies of mtDNA within a cell or an individual.^4–6^ Multiple heteroplasmies are common and widespread within the human population,^7, 8^ and more likely to be deleterious mutations and located at known disease-associated loci.^9–11^ Owing to maternal inheritance, heteroplasmic mutations are expected to be present throughout maternal lineages.^12^ Two earlier studies found that the same heteroplasmic mutations displayed varying mutant-to-wildtype allele frequencies across a wide range of tissues^13^ and in human colonic crypt stem cells^14^ in the same individuals, indicating that the observed heteroplasmic mutations must have occurred very early during embryonic development, prior to the differentiation events. Further evidence of maternal transmission of heteroplasmic mutations comes from Ding et al, who reported that approximately 30% of heteroplasmic mutations were observed in both mothers and offspring in 333 mother-offspring pairs in a low-pass whole genome sequencing (WGS) study.^9^ Similarly, a recent study by Wei et al, compared heteroplasmic mutations in 1526 mother-offspring pairs and found that about 20% of heteroplasmic mutations were concordant.^15^ Despite evidence for the inheritance of heteroplasmic mutations, the mechanisms by which heteroplasmic mutations are transmitted and maintained in humans remains to be fully elucidated.

We carried out a deep WGS study to investigate the transmission and maintenance of heteroplasmic mutations in extended pedigrees of European and African ancestry for the following (**Supplemental Table 1**). The main aim of this study was to investigate the transmission (or concordance) of heteroplasmies in first-degree and more distantly related maternal pairs in large pedigrees. We also investigated the potential functional impact of the concordant and discordant heteroplasmic mutations between paired individuals. The second aim was to investigate whether genetic loci in the nuclear genome (nDNA) are associated with heteroplasmic mutation burden in order to determine the role of nDNA in promoting or maintaining heteroplasmic mutations. To achieve these aims, we developed a comprehensive calling and quality control procedure for identifying mtDNA mutations in WGS (**Supplemental Figure 1**).^16^

## METHODS

### Study design and study participants

This study was conducted in three studies with family structures, the Framingham Heart Study (FHS), the Jackson Heart Study (JHS), and the Cleveland Family Study (CFS) (**Supplemental Tables 1 and 2**). All statistical analyses were conducted in the FHS and JHS owing to their larger sample sizes. The FHS and JHS served as a validation cohort to each other. We used CFS to further validate inconsistent results between FHS and JHS. All study participants provided written informed consent for genetic studies. All study protocols were approved by the respective Institutional Review Board of the Boston University Medical Center, the University of Mississippi Medical Center, and the Mass General Brigham (previously Partners HealthCare).

The FHS is a single-site, community-based, prospective study with extended pedigrees from three generations of European American (EA) participants.^17–19^ All FHS participants have undergone regular health examinations to collect socio-demographic characteristics and cardiovascular disease risk factors. WGS was performed in whole blood samples from 4,196 FHS participants through the Trans-Omics for Precision Medicine (TOPMed) program supported by National Heart, Lung, and Blood Institute’s (NHLBI). This study used 4,036 sequences (54% women, mean age 60) at Freeze 8 after extensive quality control (QC) procedures. The JHS is a prospective, epidemiologic investigation of CVD in African American (AA) participants from Jackson, Mississippi ^20^ across three exams beginning in 2000-2004. This study used 3,404 sequences (62% women, mean age 56) at Freeze 8 after QC. The JHS included nested pedigrees. The CFS is the largest family-based study of sleep apnea world-wide, recruiting families with a proband with diagnosed sleep apnea and matched neighborhood controls, to study sleep-disordered breathing (SDB) and cardiovascular risk.^21^ The CFS consists of 2284 individuals (46% African American) from 361 families studied on up to 4 occasions over a period of 16 years.^21^ The current study included 1,250 CFS participants (55% women, mean age 39, and 52% AA) with WGS. Based on family structures, we defined maternal lineages in both EA and AA participants. In brief, a maternal lineage contained a female founder, her offspring, and all of the grandchildren of the daughters of the founder females.^10, 22^

### Whole genome sequencing

Whole blood derived DNA was used for WGS in all participants. Data acquisition, DNA library construction, and data processing methods are described in details elsewhere (https://www.nhlbiwgs.org/topmed-whole-genome-sequencing-methods-freeze-8). Briefly, ~39X whole genome sequencing was performed at different sequencing centers: New York Genome Center, Broad Institute of MIT and Harvard, University of Washington Northwest Genomics Center, and Illumina Genomic Services.^16^ All samples for a given study were sequenced at the same center. One parent-offspring trio in the FHS was sequenced at each of four sequencing centers for QC purpose. The sequencing reads were aligned to human genome build GRCh37 at each center using similar, but not identical, processing pipelines. The resulting BAM files were transferred from each center to the TOPMed Informatics Research Center (IRC), where they were re-aligned to build GRCh37 using a common pipeline to produce a set of ‘harmonized’ BAM files. Except for the three individuals in the trio, the remaining FHS participants were sequenced at the Broad Institute of MIT and Harvard. The mean coverages were different (between 1450 and 2650) from the four repeated sequencing samples of the same parent-offspring trios by four centers, clearly showing fluctuations in sequencing manipulations across the centers (**Supplemental Table 3**). The JHS participants were sequenced at University of Washington Northwest Genomics Center, and the CFS participants were sequenced at University of Washington Northwest Genomics Center.

### Thresholds and QC procedures to identify mtDNA heteroplasmic mutations

We removed participants without the information for year at blood draw. At each mtDNA locus, we first compared an allele in sequencing data to the revised Cambridge Reference Sequence (rCRS).^23, 24^ An alternative allele in an individual refers to a different allele observed in sequencing reads when compared to the reference allele at the same locus. The program *mitoCaller* of the *mitoAnalyzer* software^9^ was applied to mtDNA sequence to derive alternative allele fractions (AAFs) for all sites to identify sequence variations including mtDNA homoplasmic variants and heteroplasmic mutations.

The Reconstructed Sapiens Reference Sequence (RSRS) was composed using a global sampling of modern human samples and samples from ancient hominids.^25^ Because the identification of alternative alleles depends on the reference sequence, we also applied RSRS to participants of both European (FHS) and African (JHS) to identify any bias in comparing results between participants of both European and African. Results in the main text were based on rCRS. All results based on RSRS were displayed in **Supplemental Materials**.

### Thresholds and QC procedures to identify mtDNA heteroplasmic mutations

In TOPMed, four sequencing centers performed sequencing with the same sequencing technology and minor fluctuations in sequencing reads exist across the sequencing centers. We applied four thresholds (t_1_ and t_2_), 1% and 99%, 2% and 98%, 3% and 97%, and 4% and 96%, to AAFs to identify the appropriate cutoffs to identify mtDNA sequence variations based on repeated mtDNA genomes of the one parent-offspring trio in the FHS from the four sequencing centers (**Supplemental Table 3**). A site was defined as a heteroplasmic mutation if its AAF was between t_1_ and t_2_ (i.e. t_1_<AAF< t_2_). A site was considered a homoplasmic variant of an alternative allele if AAF≥t_2_.

We developed a comprehensive strategy for QC of mtDNA sequence variations (**Supplemental Figure 2**). This strategy included both standard procedures used for the QC of nDNA sequence variations and procedures specific to mtDNA sequence variations. First, we investigated sequence coverage (reads) across the 16,569 mtDNA loci in the same individuals and across all individuals at each mtDNA locus. Individuals were red-flagged if their mean coverage was <500. We set an mtDNA locus as missing if the coverage was <250-fold.^9^ Second, we compared homoplasmic alleles called by genotyping arrays to the ones derived from TOPMed WGS in the same individuals.^10^ Individuals with >two inconsistent homoplasmic alleles (out of ~200 mtDNA variants genotyped by the arrays) were red-flagged and discrepancies examined. Third, we counted the number of mutations with AAFs between 25% and 75%. Based on our previous investigations, most heteroplasmic mutations displayed low AAF range (<25%).^10^ Empirically, samples having >5-10 heteroplasmic mutations whose AAFs were between 25-75% were indicative of DNA quality issues and hence were removed from subsequent analyses. Fourth, we compared the homoplasmic variants within maternal lineage members in both EA and AA participants. The nuclear mitochondrial DNA segments (NUMTs) are sequences in nDNA that show high sequence similarity to mtDNA regions.^26^ complicating the sequencing analysis of mtDNA mutations. We followed the instructions by the GATK mitochondrial pipeline to remap mtDNA sequences to the rCRS and used bedtools the ‘intersect’ option to remove NUMT regions from mtDNA bam files. We also removed several sites listed in ‘blacklisted sites’ (301,302,310, 316, 3107, and 16182 mtDNA loci) recommended by GATK (https://console.cloud.google.com/storage/browser/gatk-best-practices/mitochondria-pipeline/).

### mtDNA haplogroups

We applied Haplogrep 2 to classify mtDNA haplogroup^27–30^ based on homoplasmic mutations defined by applying the AAF>t_2_ threshold (**Supplemental Materials**).

### Concordance rate of heteroplasmies between paired individuals

To investigate transmission of heteroplasmic mutations, we formed different types of maternal pairs and random pairs in both EA and AA participants (**Supplemental Table 4**) based on maternal lineage and mtDNA haplogroup information. The maternal pairs included mother-offspring pairs, sibling-sibling pairs, and distantly related maternal pairs (i.e., grandmother-grandchild, aunt-nephew/niece, and cousin pairs on mothers’ side). For comparison purposes, we identified father-offspring pairs and also formed two types of random pairs (n=2000 each) of individuals of independent maternal lineages. The first type of random pairs belonged to the H or L mtDNA haplogroup (the largest mtDNA haplogroup in EA and AA, respectively), and the second type of random pairs belonged to different mtDNA haplogroups. We calculated the heteroplasmy concordance rate (HCR) between the paired individuals using formula (1) in EA and AA separately.

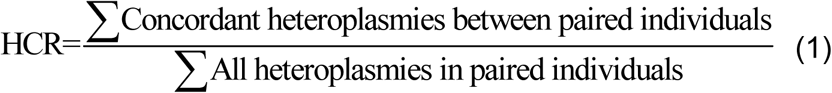

### Annotation of mtDNA heteroplasmic mutations

MitoMap was used to collect information regarding loci, regulatory elements, and previously associated phenotypes for all mtDNA variants (www.mitomap.org). For mtDNA variants within peptide-encoding genes that were non-synonymous, predicted functional effects were collected from MitImpact, a database that has compiled functional predictions across 14 bioinformatics platforms and five meta-predictors.^31^

To investigate whether heteroplasmic mutations, particularly those mutations predicted to be deleterious, were enriched in older individuals, we compared the proportions of heteroplasmic mutations in different functional categories (i.e., coding regions, nonsynonymous, and deleterious) in three age categories, 20-44, 45-64 and 65+. A nonsynonymous mutation was categorized to be deleterious if the scaled Combined Annotation-Dependent Depletion (CADD)^32^ score was 15 or above. CADD is an integrative annotation program built around more than 60 genomic features, combined with a machine-learning model trained on a binary distinction between simulated *de novo* mutations and mutations that exist in human populations. A scaled CADD score of 15 or greater indicates a mutation is amongst the top 5% of deleterious variants in the human genome.^32^ To further understand heteroplasmy inheritance, we compared proportions of heteroplasmic mutations in different functional categories among the concordant and discordant heteroplasmies in mother-offspring, sibling-sibling, and grandmother-grandchild pairs.

### Genome-wide association testing of the overall heteroplasmic burden

We quantified heteroplasmic burden by 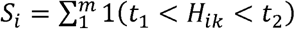. Here *H_ik_ = AAF_k_*, the alternate allele fraction at mtDNA locus *k* in the *i^th^* individual. The overall heteroplasmic burden was log-transformed and regressed on sex, age at blood drawn, batch (year at blood drawn) to obtain residuals. The inverse-normalized residuals were used as dependent variables in genome-wide testing (GWAS) with nDNA single nucleotide polymorphisms (SNPs) of minor allele frequency (MAF)>0.1% (TOPMed Freeze 8, released in February 2019, GRCH38). The Saige Linear Mixed Model was used to account for relatedness with the ENCORE server (https://encore.sph.umich.edu/). The threshold p<5×10^-8^ was used for significance to report nDNA loci associated with mtDNA heteroplasmic burden. We also included imputed cell counts (platelet, total white blood cell counts, lymphocyte or neutrophil proportions, eosinophil and basophil proportions) as additional variables in the model to obtain mtDNA CN residuals for GWAS as sensitivity analysis. Several nDNA-encoded proteins for mtDNA replication include twinkle mtDNA helicase gene^33^ (TWNK, 10q24.31), DNA polymerase subunit γ^34^ (POLG1, 15q26.1) ^34^, and the mitochondrial transcription factor gene ^35^ (TRAM, 10q21.1) are essential in the replication and maintenance of mtDNA. We searched whether any variants in these candidate genes were associated with heteroplasmic mutation burden. We also queried whether top mtDNA heteroplasmic mutation-associated loci were quantitative loci for gene expression using the eQTL database in FHS.^36^

## RESULTS

The average coverage was 2523 in FHS, 2230 in JHS, and 3000 in CFS. We determined that the 3%-97% AAF threshold gave rise to consistent numbers of homoplasmic and heteroplasmic in the same individuals from the four sequencing centers that yielded different average coverages after applying the sequential AAF thresholds to the repeated sequences of the parent-offspring trios in the FHS (Supplemental Materials, **Supplemental Table 3**), indicating that different coverage was not correlated with higher number of heteroplasmic mutations at the 3%-97% AAF threshold. Therefore, all subsequent analyses were based on 3%-97% AAF threshold. We focused on results from the FHS and JHS cohorts and used the CFS EA and CFS AA to further validate inconsistent findings between FHS and JHS. For simplicity, we used EA and AA to refer to the FHS and JHS cohort, respectively.

### Characteristics of study participants

After quality control, 4036 (FHS), 3404 (JHS) and 1250 (CFS) participants remained in the study. The FHS (mean age 60, 54% women, 100% EA) and JHS (mean age 56, 60% women, 100% AA) consisted mostly of middle aged or older participants, while the CFS (mean age 39, 55% women, 51% AA) consisted mostly of younger adults (**Supplemental Table 1**). We formed paired individuals in each cohorts based on family and maternal lineage (Supplemental materials and Supplemental Tables 2 and 4). In EA participants, we had a total of 1,545 mother-offspring pairs, 1,174 sibling-sibling pairs, 2,850 distantly related maternal pairs, and 1,119 father-offspring pairs. In AA participants, we had a total of 651 mother-offspring pairs, 689 sibling-sibling pairs, and 1,828 distantly related maternal pairs, and 263 father-offspring pairs. In each of the EA and AA cohorts, we formed 2000 pairs of unrelated individuals of the same haplogroup or different haplogroups for comparison purposes.

### Heteroplasmic mutations are ubiquitous across the mitochondrial genome

Details for identifying and quality control of heteroplasmic loci are presented in Supplemental Materials. The heteroplasmic loci were located across the entire mtDNA (Figure 1A and **Supplemental Figure 3**) and displayed similar distributions between EA and AA participants across the 37 gene and D-loop regions (Supplemental Tables 5-8). About one third of heteroplasmic mutations were located in noncoding regions and two thirds located in coding regions, and the proportion of heteroplasmic mutations in noncoding regions was much higher than expected (i.e., ~24% loci are non-coding) (p<2e-16); half of the heteroplasmic mutations in coding regions were nonsynonymous (Figure 1B). Approximately 98% of all observed heteroplasmies were present in <1% of individuals (Figure 1A). Most of the heteroplasmic mutations displayed low AAFs between 3% and 15% within the same individuals (Figure 1C), which is consistent with previous findings from our group and others.^7, 9, 10, 37, 38^ Comparison of heteroplasmic mutations identified with rCRS and RSRS, 99.6% heteroplasmic loci in EA and 96.9% loci in AA were concordant (i.e., the same number of mutations at the same mtDNA loci) in the same individuals in both EA and AA (**Supplemental Materials**, **Supplemental Table 9**).

**Figure 1.**
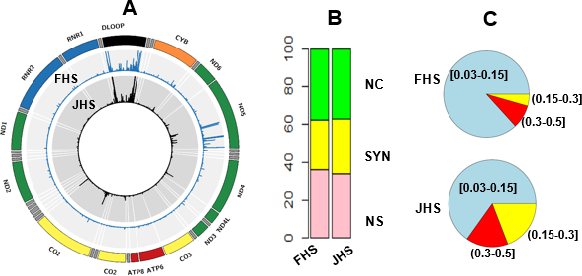
Description of mtDNA heteroplasmic mutations in European Americans (FHS) and Africans Americans (JHS). **A**, Distribution of mutations in mtDNA and number of individuals carrying mutations; **B**, Proportion of nonsynonymous (NS), synonymous (SYN) and non-coding (NC) in mutations.

### Concordant heteroplasmic mutations are observed in all types of maternal pairs

The average concordance rate of mtDNA heteroplasmic mutations referred to the proportion of mutations that were present in both of paired individuals over all heteroplasmic mutations in all paired individuals. Mother-offspring pairs displayed the highest average concordance rate, followed by sibling-sibling pairs and other pairs with more distantly maternal pairs in both EA and AA (Figure 2A and **Supplemental Figure 4**). We further found that the concordant heteroplasmic mutations displayed high correlation (R^2^~0.80) in their AAFs between paired individuals in both EA and AA (Figures 2B and 2C, **Supplemental Figures 5 and 6**). Many concordant heteroplasmies did not differ in their AAFs with advancing age (the mean age difference ~23 years between mothers and offspring pairs when blood was drawn in both EA and AA).

**Figure 2.**
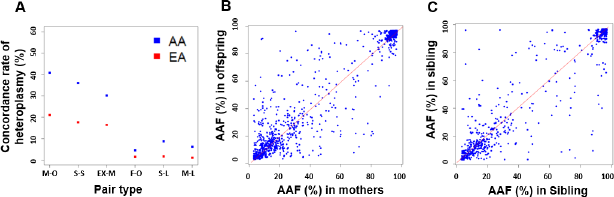
Concordant heteroplasmic mutations in paired individuals of European and African Americans. A. Concordance rate, B. Comparison of alternative allele frequency between mother-offspring pairs, C. Comparison of alternative allele frequency between sibling-sibling pairs. M-O, mother-offspring; S-S, sibling-sibling; EX-M, distantly related maternal pairs; F-O, father-offspring; S-L, unrelated pairs in the same mtDNA haplogroup; M-L, unrelated pairs of mixed mtDNA haplogroups.

As expected, father-offspring pairs and two types of unrelated pairs displayed much lower average concordance rates compared to any maternal pairs. Unrelated pairs of the same mtDNA haplogroup displayed slightly higher concordance rate than unrelated pairs of mixed haplogroups. Compared to that in unrelated pairs from mixed haplogroups, father-offspring pairs displayed a similar level of concordance rate, indicating that father-offspring transmission is unlikely to occur (Figure 2A and **Supplemental Figure 4**). Of notes, AA pairs (JHS) displayed higher concordance rates of heteroplasmic mutations compared to the respective EA pairs (FHS) when both rCRS and RSRS were applied to identify mutations; in addition, the rates were similar between the two reference sequences (**Supplemental Table 10**). The analyses in the CFS EA and CFS AA cohort further validated this finding (**Supplemental Figure 4**). Using unrelated pairs of mixed haplogroups (i.e., as a population reference), the average concordance rate was 1.5% in EA compared to 4.5% in AA (Figure 2A).

### Discordant heteroplasmic mutations are more likely to be nonsynonymous and deleterious than concordant heteroplasmic mutations

The average discordance rate of heteroplasmic mutations refers to the proportion of loci that were observed in one individual but not another in a pair over all heteroplasmic mutation observed in paired individuals. A larger proportion of discordant heteroplasmies were nonsynonymous compared to concordant heteroplasmic mutations (Figure 3). For example, in sibling-sibling pairs of EA participants, 30.8% of discordant loci were nonsynonymous while 12.4% of concordant loci were nonsynonymous (p=2.2e-16). Furthermore, a larger proportion of discordant mutations were deleterious compared to concordant ones: 9.9% of discordant loci versus 5.8% of concordant loci were deleterious in EA mother-offspring pairs (p=1.2e-7); 13.9% of discordant loci versus 4.3% of concordant loci were deleterious in EA sibling-sibling pairs (p=1.9e-13) (Figure 3). Similar findings were observed in mother-offspring and sibling-sibling pairs in AA participants (**Supplemental Figure 7**).

**Figure 3.**
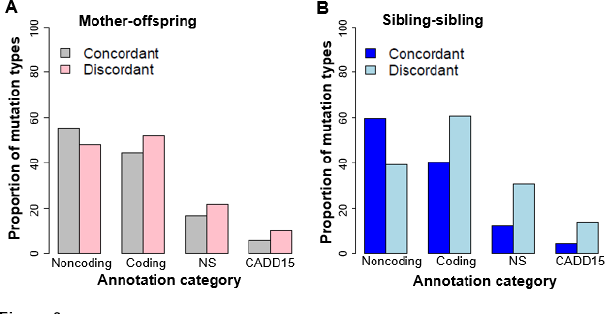
Discordant heteroplasmic mutations are more likely to be deleterious than concordant heteroplasmic mutations. The proportion of concordant heteroplasmic mutations in noncoding, coding regions, being nonsynonymous (NS), and being deleterious with CADD ≥15 (CADD15) in Mother-offspring and sibling-sibling pairs in the Framingham Heart Study (FHS).

### Older individuals carry a higher burden of deleterious heteroplasmic mutations

Of all heteroplasmic mutations in both paired individuals, older individuals carried a larger proportion of discordant mutations compared to younger individuals. For example, of all heteroplasmic mutations in EA mother-offspring pairs, 21.2% mutations were concordant between mothers and offspring, 45.5% were only observed in mothers while 33.2% were present only in offspring (p<2.2e-16). In contrast, offspring pairs carried similar proportions of discordant heteroplasmic mutations between each other because sibling-sibling pairs were formed without regard for age (Figure 4A). Similar findings were also observed in mother-offspring and offspring-offspring pairs in AA (**Supplemental Figure 8A**).

**Figure 4.**
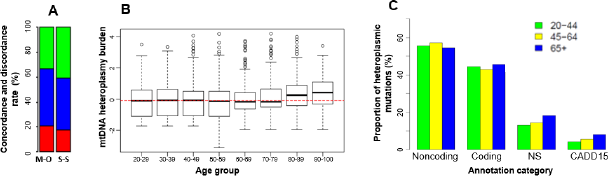
Heteroplasmy and age in participants of European descent. **A**.Concordant (red color) and discordant (green and blue colors) heteroplasmic mutations between mother-offspring (M-O) and sibling-sibling (S-S) pairs, in mother-offspring bar, the blue portion is the proportion of discordant heteroplasmic mutations in mothers only and the green portion is the proportion of discordant mutations in offspring only, the green and blue proportions in the sibling-sibling bar represent the proportion of mutations in sibling pairs; **B**. The overall burden of mtDNA heteroplasmic mutations in different age groups; **C**. The proportion of heteroplasmic mutations according to functional annotations in individuals of 20-44, 45-64 and 65 or above; Noncoding, the proportion of heteroplasmies in noncoding regions. Coding, the proportion of heteroplasmies in 13 protein-coding regions; NS, the proportion of nonsynonymous in coding heteroplasmies; CADD15, the proportion of possibly deleterious nonsynonymous mutations with CADD≥15 in coding heteroplasmies.

A comparison of the heteroplasmic burden (i.e., the total number of heteroplasmic mutations in a region or across mtDNA in an individual) in age groups found that the heteroplasmic burden was at a similar level in participants across a wide age range, but became significantly greater in individuals ≥ 80 years in EA (p<0.002) and ≥70 years in AA participants (p<0.004) (Figure 4B and **Supplemental Figure 8B**). This trend did not change after adjusting for cell counts (**Supplemental Figure 9**). We further compared heteroplasmic burden in age group, 20-44, 45-64, and ≥65 years, according to functional annotations (noncoding, coding, nonsynonymous, and possibly deleterious with CADD≥15) (Figure 4C, **Supplemental Figure 8C**). Both AA and EA participants carried a higher burden of heteroplasmic mutations in noncoding regions than in coding regions across all age groups. Participants of 65 years and older carried a greater burden of nonsynonymous heteroplasmic mutations compared to younger individuals in both EA and AA participants. The burden of deleterious heteroplasmic mutations increased with advancing age (20-44, 45-64, ≥65 years) with trend test p=0.010 in both EA and AA (Figure 4C, **Supplemental Figure 8C**). The heteroplasmic burden was not significantly different between men and women in EA (p>0.05), but was greater in men compared to women in AA (p=0.0004) after accounting for age and batch effects (**Supplemental Table 11**).

### The heteroplasmic burden is moderately related to white blood cell count and differential count, and mtDNA haplogroup

We found that the total heteroplasmic burden was modestly associated with imputed white blood cell counts and differentials. The cell counts jointly explained 1.3% (p=1e-7, EA) and 0.5% (p=0.001, AA) of the heteroplasmic burden (**Supplemental Table 12**). mtDNA haplogroups were significantly associated with the total heteroplasmic burden, explaining 5.3% (p<1e-16, EA) and 8.6% (p<1e-16, AA) of the variability in heteroplasmic burden. Pairwise comparisons demonstrated that the heteroplasmic burden differed significantly between haplogroups (**Supplemental Table 13**).

### Chromosome 11p11-12 region is significantly associated with the overall heteroplasmic mutation burden

Using the SAIGE^39^ method provided by ENCORE (The University of Michigan), 15,037,233 (Freeze 8, EA) and 29,391,078 (Freeze 8, AA) nDNA variants (MAF>0.1%) were tested for association with the overall burden of heteroplasmy. There was little or no inflation (genomic inflation factor λ=1.00 in EA and 1.05 in AA) (Supplemental Figures 10 and 11). Hundreds of variants at the Olfactory Receptor Family 4 Subfamily C Member 12 region (*OR4C12*, 11p11.12) were associated with heteroplasmic burden (at p <5e-8 in both cohorts (Supplemental Figures 10 and 11). In EA, the lowest p-value was observed at rs2773516 (p= 7.1E-10; MAF=0.40, A/C; chr11: 50427662), which explained 1.3% of variability in heteroplasmy burden. In AA, the lowest p-value was observed at rs36145545 (p= 9.4E-11; MAF=0.38, G/A; chr11:50074179), which explained 1.2% of variability in heteroplasmy burden. We performed a sensitivity analysis in the same sample size with cell counts as additional covariates. The GWAS signals attenuated in the EA (**Supplemental Figure 12**), but it remained unchanged in the AA, after adjusting for cell counts (**Supplemental Figure 13**).

Meta-analysis of variants with MAF ≥1% that were associated with heteroplasmic burden identified 1,277 SNPs at 11p11.12 with p<5e-8 (the top signal at rs779031139, p=2.0e-18; frequency of A allele =0.38 in EA and 0.67 in AA) in meta-analysis of EA and AA (**Supplemental Table 14**). The significant SNPs from meta-analysis jointly explained 3.5% (EA) and 3.6% (AA) of interindividual variation in heteroplasmic burden. We explored whether these significant SNPs were also associated with reported phenotypes in a large GWAS database (https://www.ebi.ac.uk-/gwas/). Five heteroplasmic burden-associated SNPs were associated with height^40^ and depression^41^ in previous GWAS (**Supplemental Table 15**). None of the variants in TWNK^33^, TRAM^35^, and POLG1^34^ were significantly associated with heteroplasmic mutation burden. Many of the top SNPs (n=1,031) act as long-range *cis* regulators of PTPRJ (protein tyrosine phosphatase receptor type J) (**Supplemental Table 16**). PTPRJ is a signaling molecule that regulates a variety of cellular processes, including cell growth, differentiation, mitotic cycle, and oncogenic transformation.

## DISCUSSION

Understanding the transmission of mtDNA heteroplasmic mutations is essential to study the role of mtDNA in relation to a variety of disease phenotypes. Despite maternal inheritance and the lack of germline recombination,^42, 43^ the inheritance and transmission of heteroplasmies is complex. In this study, we compared the concordance rates of heteroplasmic mutations in mother-offspring and other types of maternal pairs in EA and AA. We replicated previous reports that many heteroplasmic mutations are transmitted from mothers to offspring.^9, 15^ In addition, we observed that concordant heteroplasmic mutations were present in sibling-sibling pairs and more distantly related maternal pairs. Many of the transmitted heteroplasmic mutations exhibited similar alternative allele fractions between maternal pairs. The concordance rate of heteroplasmies was highest between the founder women and their offspring, and become lower in more distantly related pairs, and continued to decay to a low level in unrelated pairs within the same mtDNA haplogroup. Based on functional annotation, concordant heteroplasmic mutations were more likely to be non-coding and synonymous, and thus, they were less likely to impact mitochondrial function. In contrast, the discordant heteroplasmic mutations were more likely to be non-synonymous and deleterious, indicating that natural selection may play a role in constraining the transmission of pathogenic heteroplasmic alleles.

A larger proportion of heteroplasmic mutations, however, were discordant between mothers and offspring. The discordant heteroplasmies may result from combinations of the three events: a bottleneck phenomenon, clonal expansion, and *de novo* mutations. The bottleneck phenomenon refers to a condition whereby a mixture of mtDNA molecules are present in a mother’s germ cell, while a subset of mtDNA molecules are transmitted into an oocyte.^12^ Therefore, an oocyte contains only a subset of heteroplasmies of the mother. Subsequent clonal expansion (i.e., the rapid replication of the mtDNA) occurs during oocyte maturation.^44^ Owing to the bottleneck phenomenon and clonal expansion, some heteroplasmies only exist in mothers but not in offspring, and only a proportion of the heteroplasmies are concordant between siblings. After oocyte maturation and clonal expansion, some undetectable heteroplasmies in mothers may become detectable (and thus are considered *de novo*) in offspring due to the increase of alternative allele frequencies. The fact that a much larger proportion of heteroplasmies was observed in older individuals within maternal pairs may reflect both *de novo* mutations and an increase in alternative allele frequencies during aging. Further research is needed to investigate the functional significance of *de novo* heteroplasmies and those that increase their AAFs during aging.

In this cross-sectional study, we observed that the level of heteroplasmy burden in whole blood was at similar levels, on average, across most of the age ranges in participants of EA and AA. The lifespan of white blood cells ranges from 13-20 days.^45^ Old white blood cells are destroyed and replaced by newly generated white blood cells from stem cells in bone marrow. This turnover of white blood cells may explain the stable heteroplasmic burden across the most of the age range of our study sample. However, this turnover mechanism of white blood cells is not able to explain the observation of a significant increase in heteroplasmy burden in older age. It can neither explain the observation that the burden of deleterious (combined annotation-dependent depletion score ≥15) heteroplasmies was significantly correlated with advancing age (20-44, 45-64, ≥65 years). Therefore, there might exist a molecular ‘brake’ on the regulation of mtDNA heteroplasmic mutations. This molecular ‘brake’ is likely to become less effective during aging. The accumulation of deleterious mtDNA mutations disturbs the integrity of mtDNA, which may lead to impaired mitochondrial function contributing to the pathogenesis of age-related disease.^46^ The true impact of heteroplasmic mutations on human health remains to be investigated.

Because nDNA plays an essential role in the replication of mtDNA, we conducted a GWAS of heteroplasmy burden. Of note, none of variants in the candidate mtDNA regulatory genes (*TWNK*^33^, *TRAM*^35^, and *POLG1*^34^) are significantly associated with heteroplasmy burden. Instead, we found that a long chromosomal region at 11p11.12 is significantly associated with overall heteroplasmy burden. This region harbors the olfactory receptor family 4 gene and several pseudogenes that do not show clear functional inference regarding their roles in promoting or maintaining mtDNA heteroplasmy. Further exploration revealed that most of the top variants in this region are strong long-range eQTLs of *PTPRJ* (protein tyrosine phosphatase receptor type J). *PTPRJ* is involved in dephosphorylation of many important proteins in cell adhesion, migration, proliferation and differentiation. A previous study in lung cancer patients found that *PTPRJ* negatively modulated several proteins that are related to mitochondrial functions,^47^ including acyl-coenzyme A thioesterase 9, NADH dehydrogenase [ubiquinone] iron-sulfur protein 2, the mitochondrial import receptor subunit TOM70, the ATP synthase subunit gamma, the ATP synthase subunit alpha, and the carbamoyl-phosphate synthase.^47^ Further research is needed to investigate whether *PTPRJ* plays similar roles in healthy individuals.

### Strengths and limitations

We developed comprehensive quality control strategies for the identification of heteroplasmic mutations using WGS. Consistent quality control and identification of heteroplasmic mutations is critical to facilitate trait-association analysis of heteroplasmic mutations across the TOPMed cohorts. We applied the same AAF thresholds to identify heteroplasmic mutations in participants of both EA and AA. Despite the multiple strengths in this study, several limitations should be noted. We studied the characteristics of mtDNA heteroplasmies in whole blood. The location, variant allele frequency, and burden of heteroplasmic mutations in blood may not be reflective of that in other tissues. Studies of mtDNA heteroplasmies in specific tissues may reveal different information with respect to metabolic (e.g., adipose tissue), heart (e.g., cardiac muscle) and aging-related (e.g., brain) diseases. Peripheral blood is easily accessible and least invasive to obtain, it circulates through all tissues and organs, and may provide important information in mtDNA disease susceptibility.

## Disclaimer

The views expressed in this manuscript are those of the authors and do not necessarily represent the views of the National Heart, Lung, and Blood Institute; the National Institutes of Health; or the U.S. Department of Health and Human Services.

## TOPMed Acknowledgements

Molecular data for the Trans-Omics in Precision Medicine (TOPMed) program was supported by the National Heart, Lung and Blood Institute (NHLBI). See the TOPMed Omics Support Table in the Supplemental Materials for study specific omics support information. Core support including centralized genomic read mapping and genotype calling, along with variant quality metrics and filtering were provided by the TOPMed Informatics Research Center (3R01HL-117626-02S1; contract HHSN268201800002I). Core support including phenotype harmonization, data management, sample-identity QC, and general program coordination were provided by the TOPMed Data Coordinating Center (R01HL-120393; U01HL-120393; contract HHSN268201800001I). We gratefully acknowledge the studies and participants who provided biological samples and data for TOPMed.

See Supplemental Materials for additional study specific acknowledgements.

## Notes

### Competing Interest Statement

The authors have declared no competing interest.

