## Supplemental Materials for "Presence and Transmission of Mitochondrial Heteroplasmic Mutations in Human Populations of European and African Ancestry"

Acknowledgement

The Framingham Heart Study (EA) (phs000974.v1.p1) is supported and conducted in collaboration with the Broad Institute of MIT and Harvard (HHSN268201500014C, 3R01HL092577-06S1, 3U54HG003067-12S2, HHSN268201600034I, 3U54HG003067-12S2, 3R01HL092577-06S1), NWGC (HHSN268201600032I), Keck MGC (HHSN268201600038I) 3U54HG003067-12S2contracts from the National Heart, Lung, and Blood Institute (NHLBI) and the National Institute for Minority Health and Health Disparities (NIMHD). The FHS acknowledges the support of contracts NO1-HC-25195, HHSN268201500001I and 75N92019D00031 from the National Heart, Lung and Blood Institute and grant supplement R01 HL092577-06S1 for this research. We also acknowledge the dedication of the FHS study participants without whom this research would not be possible. Dr. Vasan is supported in part by the Evans Medical Foundation and the Jay and Louis Coffman Endowment from the Department of Medicine, Boston University School of Medicine. C.L. is supported by individual funding (R01AG059727). JLF is supported by a K01HL143142 and a TOPMed Junior Investigator Award (5U01HL117626-05). The authors also wish to thank the staff and participants of the EA.

The Jackson Heart Study (JHS) (phs000964.v1.p1) is supported and conducted in collaboration with NWGC (3U54HG003067-12S2) contracts from the National Heart, Lung, and Blood Institute (NHLBI), Jackson State University (HHSN268201300049C, HHSN268201300050C, HHSN268201800013I), Tougaloo College (HHSN268201300048C, HHSN268201800014I), the Mississippi State Department of Health (HHSN268201800015I), and the University of Mississippi Medical Center (HHSN268201300046C and HHSN268201300047C, HHSN268201800010I, HHSN268201800011I and HHSN268201800012I) contracts from the National Heart, Lung, and Blood Institute (NHLBI) and the National Institute for Minority Health and Health Disparities (NIMHD). WGS was performed at University of Washington (HHSN268201100037C). The authors also wish to thank the staffs and participants of the JHS. HHSN268201100037C

The Cleveland Family Study (CFS) (phs000954) is supported and conducted by NWGC (HHSN268201600032I, 3R01HL098433-05S1), and also supported in part by National Institutes of Health grants (R01-HL046380, KL2-RR024990, R35-HL135818, and R01-HL113338).

**TOPMed Omics Support Center**

| TOPMed Accession # | TOPMed Project | Parent Study | TOPMed Phase | Omics Center | Omics Support | Omics Type |
| --- | --- | --- | --- | --- | --- | --- |
| phs000954 | CFS | CFS | 3.5 | NWGC | HHSN268201600032I | WGS |
| phs000954 | CFS | CFS | 1 | NWGC | 3R01HL098433-05S1 | WGS |
| phs000974 | AFGen | FHS AFGen | 1 | Broad Genomics | 3R01HL092577-06S1 | WGS |
| phs000974 | FHS | FHS | pilot; 4.5 | NWGC | HHSN268201600032I | RNASeq |
| phs000974 | FHS | FHS | 5 | NWGC | HHSN268201600032I | RNASeq |
| phs000974 | FHS | FHS | pilot; 4.5 | Broad BIDMC Proteomics | HHSN268201600034I | Proteomics |
| phs000974 | FHS | FHS | 5 | Broad Metabolomics | HHSN268201600034I | Metabolomics |
| phs000974 | FHS | FHS | 5 | Keck MGC | HHSN268201600038I | Methylomics |
| phs000974 | FHS | FHS | 1 | Broad Genomics | 3U54HG003067-12S2 | WGS |
| phs000964 | JHS | JHS | 1 | NWGC | HHSN268201100037C | WGS |

**Identification and quality control of heteroplasmic mutations**

At each mtDNA locus, an allele in sequencing data was compared to the allele of the revised Cambridge Reference Sequence (rCRS).^22, 23^ An alternative allele in an individual refers to a different allele observed in sequencing reads when compared to the reference allele at the same locus. The program *mitoCaller* of the *mitoAnalyzer* software^9^ was applied to mtDNA sequence to derive alternative allele fractions (AAFs) for all sites to identify sequence variations including mtDNA homoplasmic variants and heteroplasmic mutations. Although the four sequencing centers used the same sequencing technology, minor fluctuations in sequencing reads exist across the sequencing centers. We applied four thresholds (t_1_ and t_2_), 1% and 99%, 2% and 98%, 3% and 97%, and 4% and 96%, to AAFs to identify the appropriate cutoffs to identify mtDNA sequence variations based on repeated mtDNA genomes of the one parent-offspring trio in the FHS from the four sequencing centers. A site was defined as a heteroplasmic mutation if its AAF was between t_1_ and t_2_ (i.e. t_1_<AAF< t_2_). A site was considered a homoplasmic variant of an alternative allele if AAF≥t_2_.

We developed a comprehensive strategy for QC of mtDNA sequence variations (**Supplemental** **Figure 2**). This strategy included both standard procedures used for the QC of nDNA sequence variations and procedures specific to mtDNA sequence variations. First, we investigated sequence coverage (reads) across the 16,569 mtDNA loci in the same individuals and across all individuals at each mtDNA locus. Individuals were red-flagged if their mean coverage was <500. We set an mtDNA locus as missing if the coverage was <50-fold.^9^ Second, we compared homoplasmic alleles called by genotyping arrays to the ones derived from TOPMed WGS in the same individuals.^10^ Individuals with >two inconsistent homoplasmic alleles (out of ~200 variants genotyped by the arrays) were red-flagged and discrepancies examined. Third, we counted the number of mutations with AAFs between 25% and 75%. Based on our previous investigations, most heteroplasmic mutations displayed low AAF range (<25%).^10^ Empirically, samples having >5-10 heteroplasmic mutations whose AAFs were between 25-75% were indicative of DNA quality issues and hence were removed from subsequent analyses. Fourth, we compared the homoplasmic variants within maternal lineage members in both EA and AA participants. The nuclear mitochondrial DNA segments (NUMTs) are sequences in nDNA that show high sequence similarity to mtDNA regions.^24^ complicating the sequencing analysis of mtDNA mutations. We followed the instructions by the GATK mitochondrial pipeline to remap mtDNA sequences to the rCRS and used bedtools the ‘intersect’ option to remove NUMT regions from mtDNA bam files. We also removed several sites listed in ‘blacklisted sites’ (301,302,310, 316, 3107, and 16182 mtDNA loci) recommended by GATK (<https://console.cloud.google.com/storage/browser/gatk-best-practices/mitochondria-pipeline/>).

We applied consistent QC procedures to all participants. mtDNA sequences were extracted in 4076 FHS individuals in FHS. After QC, 41 individuals were excluded due to 1) wrong maternal relationship (n=2); 2) possible low DNA quality identified by mtDNA specific procedures and comparison with previous genotyping data (n=7); 3) bad metrics by nuclear DNA (nDNA) QC procedures; 4) repeated samples (n=9); 5) and unmatched blood draw dates on any exams (n=17). mtDNA is maternally inherited. We counted homoplasmic mutations based on a relaxed threshold 25%-75%. We applied the same quality control procedures to JHS and CFS samples. We did not find individuals with wrong maternal relationship. There was no repeated samples. The three cohorts were sequenced in two different genotyping centers. The FHS and JHS included middle aged participants and CFS included younger participants when blood was drawn for sequencing (**Supplemental Table 1**).

**Threshold to identify heteroplasmic mutations**

To select the most appropriate threshold to define mtDNA mutations, we applied several AAF thresholds to the repeated sequences of the parent-offspring trio individuals. The 1%-99% threshold gave rise to more sequence variants and corresponding homoplasmy and heteroplasmy calls in repeated samples from the same individuals of. Higher coverage in a sample, however, did not give rise to increased numbers of heteroplasmic mutations at the 1%-99% threshold. Both the 3%-97% and 4%-96% thresholds yielded consistent numbers of homoplasmic variants and heteroplasmic mutations in repeated samples from the same individuals regardless of sequence coverage/sequence centers (**Supplemental** **Table 3**). In addition, the mother-offspring pairs showed consistent numbers of homoplasmic variants at the 3%-97% threshold in both the FHS and JHS. Therefore, we chose the 3%-97% threshold to define sequence variants. A site was deemed as heteroplasmic if its AAF was between 3% and 97% in an individual. A site was deemed as homoplasmic if its AAF was ≥97% in an individual.

With the 3%-97% AAF threshold and extensive QC procedures, we identified 3411 loci with variations and of them 2,262 loci were heteroplasmic in at least one EA (FHS) participant (**Supplemental Table 6).** We identified 3508 loci with variations and of them 1,643 loci were heteroplasmic in at least one AA (JHS) participants (**Supplemental Table 7)**. In CFS EA, we identified a total of 1,640 variation loci, of them, 348 are heteroplasmic mutations (**Supplemental Table 8**). In CFS AA, we identified a total of 1,618 variation loci, of them 334 are heteroplasmic mutations (**Supplemental Table 7**). Of notes, several heteroplasmic loci contained two or more alleles, and most of these loci are located in the D-loop region (**Supplemental Tables 6-8).**

**Comparison of heteroplasmic mutations in participants in the Framingham Heart Study and Jackson Heart Study**

We compared the identified mtDNA heteroplasmic loci and the number of loci carried by individuals based on rCRS and RSRS. In FHS, the rCRS and RSRS identified exactly the same loci (n=2,262), and only 9 loci distributed differently in the FHS participants (**Supplemental Table 9**). As expected, most individuals carried the same number of heteroplasmic mutations at most loci except for 9 loci. The Pearson correlation of the number of heteroplasmic mutations across individuals was 0.998. In JHS participants, rCRS identified 1,643 heteroplasmic mutations and RSRS identified 1,645 heteroplasmic mutations, with 1,642 loci overlapped between the two references (**Supplemental Table 9**). In addition to 6 non-overlapped loci, 51 overlapped loci distributed differently in JHS. Most of these 51 loci (80%) distributed differently by only one individual in JHS (**Supplemental Table 9)**. The 16189 loci, located in the hypervariable, non-coding region distributed most differently in both FHS and JHS.

**Pedigree structure and moternal lineages in FHS and JHS**

We used Freeze 8 whole genome sequencing data in all participating cohorts. The FHS contained extended pedigrees while JHS contained nested families. The CFS consisted of family structures by design. The CFS is the largest family-based study of sleep apnea world-wide, recruiting families with a proband with diagnosed sleep apnea with matching neighborhood controls (**Supplemental Table 2a**). A maternal lineage contained a women founder, her daughters and sons, and all the grandchildren of the daughters of the founder women. Moternal lineages were assigned to founder mothers and passed down to their daughters and sons, and all the grandchildren of the daughters of the founder women. We formed maternal lineages in all participants using their family structures (**Supplemental Table 2b**).

**mtDNA haplogroup classification**

The major mtDNA haplogroups are similar between the two EA populations. For examples, the H haplogroup is the largest one, followed by U, J, T, and K in EA population (**Supplemental Table 13A**). Similarly, the largest mtDNA haplogroup was L3, followed by L2 and L1 haplogroups in African Americans (**Supplemental Table 13B**).

**Formation of paired individuals**

To investigate concordance rates, we formed different types of maternal pairs and random pairs in both EA and AA participants based on maternal lineage and mtDNA haplogroup information (**Supplemental Table 4)**. The maternal pairs included mother-offspring pairs, sibling-sibling pairs, and distantly related maternal pairs (i.e., grandmother-grandchild, aunt-nephew/niece, and cousin pairs on mothers’ side). The FHS contained 1311 mother-offspring pairs, 1031 sibling-sibling pairs, and 2376 more distant maternal pairs (including cousin pairs, grandmother-grandchildren, and aunt-niece/nephew pairs on mothers’ side). To compare concordance and discordant heteroplasmic mutations, we formed three additional pairs in the FHS, including 929 father-offspring, and 2000 pairs of unrelated individuals of the same mtDNA H haplogroup (the largest haplogroup in EA) or mixed haplogroups (**Supplemental Table 4**). We formed the same types of pairs in the JHS, including 347 mother-offspring pairs, 640 sibling-sibling pairs, 1013 other maternal pairs, 139 father-offspring pairs, 2000 pairs of unrelated individuals of the same mtDNA L3 haplogroup (the largest haplogroup in AA) or mixed haplogroups. In EA participants in the CFS cohort, we formed 247 mother-offspring pairs, 151 offspring-offspring pairs, 889 other maternal pairs, 190 father-offspring pairs, and 560 pairs of unrelated individuals of mixed mtDNA haplogroups. In AA participants in the CFS cohort, we formed 289 mother-offspring pairs, 60 offspring-offspring pairs, 1,178 other maternal pairs, 125 father-offspring pairs, and 91 pairs of unrelated individuals of mixed mtDNA haplogroups (**Supplemental Table 4**).

**Supplemental Table 1**. Participant characteristics in Framingham Heart Study (FHS), Jackson Heart Study (JHS), and Cleveland Family Study (CFS).

| Traits | FHS (EA)  (n=4,036) | JHS (AA)  (n=3,406) | CFS (EA)  (n=611) | CFS (AA)  (n=639) |
| --- | --- | --- | --- | --- |
| Women | 2159(54.6%) | 2142 (62.9%) | 321 (52.5%) | 363 (57.6) |
| Age | 59.9±15.5 | 55.7±12.8 | 41.4 (19.0) | 38.5 (19.4) |
| Sequencing center | Broad | UW | UW | UW |
| Median coverage | 2523 | 2230 | 2998 | 3001 |

Data are presented as mean ± SD or n %. EA, European American; AA, African American; Broad, Broad Institute of MIT and Harvard; UW, University of Washington Northwest Genomics Center.

**Supplemental Table 2a**. The size and number of pedigrees in Framingham Heart Study (FHS), Jackson Heart Study (JHS), and Cleveland Family Study (CFS).

| **FHS (n=4036)** | | **JHS (n=3404)** | | **CFS (n=1250)** | | | |
| --- | --- | --- | --- | --- | --- | --- | --- |
| **EA** | | **AA** | | **EA (n=611)** | | **AA (n=639)** | |
| **size** | **N of families** | **Size** | **N of families** | **Size** | **N of families** | **Size** | **N of families** |
| 1 | 1 | 1 | 1914 | 1 | 27 | 1 | 13 |
| 2 | 13 | 2 | 191 | 2 | 3 | 2 | 2 |
| 3 | 116 | 3 | 64 | 3 | 3 | 3 | 3 |
| 4 | 57 | 4 | 35 | 4 | 1 | 4 | 4 |
| 5 | 47 | 5 | 20 | 5 | 2 | 5 | 1 |
| 6 | 41 | 6 | 19 | 6 | 2 | 6 | 28 |
| 7 | 39 | 7 | 13 | 7 | 1 | 7 | 21 |
| 8 | 19 | 8 | 8 | 8 | 1 | 8 | 1 |
| 9 | 30 | 9 | 7 | 9 | 14 | 9 | 28 |
| 10 | 14 | 10 | 3 | 10 | 23 | 10 | 13 |
| 11 | 13 | 11 | 2 | 11 | 17 | 11 | 8 |
| 12 | 9 | 12 | 4 | 12 | 12 | 12 | 6 |
| 13 | 12 | 14 | 1 | 13 | 6 | 13 | 7 |
| 14 | 3 | 16 | 2 | 14 | 7 | 14 | 3 |
| 15 | 4 | 17 | 1 | 15 | 7 |  |  |
| 16 | 5 | 20 | 1 | 16 | 5 |  |  |
| 17 | 5 | 21 | 1 |  |  |  |  |
| 18 | 4 | 22 | 1 |  |  |  |  |
| 19 | 4 | 23 | 3 |  |  |  |  |
| 20 | 6 | 24 | 1 |  |  |  |  |
| 21 | 3 | 25 | 1 |  |  |  |  |
| 22 | 2 |  |  |  |  |  |  |
| 23 | 1 |  |  |  |  |  |  |
| 26 | 2 |  |  |  |  |  |  |
| 27 | 1 |  |  |  |  |  |  |
| 29 | 1 |  |  |  |  |  |  |
| 32 | 1 |  |  |  |  |  |  |
| 34 | 1 |  |  |  |  |  |  |
| 40 | 1 |  |  |  |  |  |  |
| 41 | 1 |  |  |  |  |  |  |
| 42 | 1 |  |  |  |  |  |  |
| 44 | 1 |  |  |  |  |  |  |
| 45 | 1 |  |  |  |  |  |  |
| 60 | 1 |  |  |  |  |  |  |
| 97 | 1 |  |  |  |  |  |  |
| 117 | 1 |  |  |  |  |  |  |
| 150 | 1 |  |  |  |  |  |  |
| 235 | 1 |  |  |  |  |  |  |

**Supplemental Table 2b**. The size and number of moternal lineages in Framingham Heart Study (FHS) and Jackson Heart Study (JHS), and Cleveland Family Study (CFS).

| **Size of maternal  lineage** | **Number of the lineage** | | | |
| --- | --- | --- | --- | --- |
|  | **FHS**  **(EA)** | **JHS**  **(AA)** | **CFS**  **(EA)** | **CFS**  **(AA)** |
| 1 | 661 | 2095 | 140 | 94 |
| 2 | 680 | 239 | 3 | 1 |
| 3 | 194 | 62 | 2 | 2 |
| 4 | 116 | 45 | 55 | 1 |
| 5 | 82 | 26 | 39 | 58 |
| 6 | 31 | 16 | 16 | 28 |
| 7 | 18 | 6 | 5 | 24 |
| 8 | 7 | 9 | 6 | 11 |
| 9 | 8 | 4 | 4 | 9 |
| 10 | 7 | 3 | 2 | 4 |
| 11 | 2 | 2 | 1 | 2 |
| 12 | 1 | 1 |  | 4 |
| 13 | 0 | 1 |  |  |
| 14 | 0 | 1 |  |  |
| 15 | 1 | 0 |  |  |

**Supplemental Table 3**. Framingham Heart Study repeated samples by four sequencing centers in TOPMed

| **TOPMed ID** | | **Mean coverage** | **Sequencing  center** | **1% and 99%** | | **2% and 98%** | | **3% and 97%** | | **4% and 96%** | |
| --- | --- | --- | --- | --- | --- | --- | --- | --- | --- | --- | --- |
|  |  |  |  | **Het** | **Hom** | **Het** | **Hom** | **Het** | **Hom** | **Het** | **Hom** |
| IND 1 | NWD321439 | 2665 | Broad | 6 | 16 | 4 | 16 | 2 | 16 | 2 | 16 |
| IND 1 | NWD465832 | 2057 | Illumina | 9 | 16 | 4 | 16 | 2 | 16 | 2 | 16 |
| IND 1 | NWD433184 | 1675 | NYGC | 57 | 11 | 6 | 15 | 2 | 16 | 2 | 16 |
| IND 1 | NWD481739 | 1592 | UW | 13 | 16 | 3 | 16 | 2 | 16 | 2 | 16 |
| IND 2 | NWD813793 | 1846 | Broad | 9 | 23 | 4 | 25 | 3 | 25 | 3 | 25 |
| IND 2 | NWD507659 | 2333 | Illumina | 9 | 24 | 4 | 24 | 3 | 24 | 3 | 24 |
| IND 2 | NWD143123 | 1848 | NYGC | 10 | 25 | 4 | 25 | 3 | 25 | 3 | 25 |
| IND 2 | NWD284313 | 1844 | UW | 11 | 25 | 5 | 25 | 3 | 25 | 3 | 25 |
| IND 3 | NWD985536 | 1458 | Broad | 13 | 12 | 3 | 16 | 3 | 16 | 2 | 16 |
| IND 3 | NWD820361 | 2146 | Illumina | 16 | 14 | 5 | 16 | 3 | 16 | 3 | 16 |
| IND 3 | NWD808124 | 1694 | NYGC | 16 | 12 | 3 | 16 | 3 | 16 | 3 | 16 |
| IND 3 | NWD641266 | 1937 | UW | 17 | 14 | 3 | 16 | 3 | 16 | 3 | 16 |

**Supplemental Table 4**. Paired individuals in European and African American participants

| **Pair type** | **EA** | | **AA** | |
| --- | --- | --- | --- | --- |
|  | FHS | CFS EA | JHS | CFS AA |
| Mother-offspring pairs | 1311 | 234 | 374 | 277 |
| Sibling-sibling pairs | 1031 | 143 | 640 | 49 |
| Other maternal pairs | 2376 | 474 | 1013 | 815 |
| ^1^Unrelated pairs in H or L3 mtDNA haplogroup | 2000 | 2000 | 2000 | 2000 |
| ^2^Unrelated pairs of mixed mtDNA haplogroups | 2000 | 2000 | 2000 | 2000 |
| Father-offspring pairs | 929 | 190 | 139 | 124 |

EA, European American; AA, African American

**Supplemental Table 5**. Distribution of heteroplasmic mutations in the Framingham Heart Study and Jackson Heart Study

| **Location** | **Number of heteroplasmic mutations** | | **Frequency of heteroplasmic mutations** | |
| --- | --- | --- | --- | --- |
|  | **FHS** | **JHS** | **FHS** | **JHS** |
| D-loop | 286 | 226 | 0.130474 | 0.143564 |
| MT-ATP6 | 107 | 77 | 0.047811 | 0.047649 |
| MT-ATP8 | 34 | 27 | 0.015192 | 0.016708 |
| MT-CO1 | 203 | 166 | 0.090706 | 0.102723 |
| MT-CO2 | 78 | 65 | 0.034853 | 0.040223 |
| MT-CO3 | 90 | 82 | 0.040214 | 0.050743 |
| MT-CYB | 174 | 134 | 0.077748 | 0.082921 |
| MT-ND1 | 98 | 80 | 0.043789 | 0.049505 |
| MT-ND2 | 130 | 77 | 0.058088 | 0.047649 |
| MT-ND3 | 33 | 28 | 0.014745 | 0.017327 |
| MT-ND4 | 125 | 88 | 0.055853 | 0.054455 |
| MT-ND4L | 27 | 18 | 0.012064 | 0.011139 |
| MT-ND5 | 260 | 177 | 0.116175 | 0.10953 |
| MT-ND6 | 67 | 47 | 0.029937 | 0.029084 |
| MT-RNR1 | 147 | 62 | 0.065684 | 0.038366 |
| MT-RNR2 | 205 | 126 | 0.0916 | 0.07797 |
| MT-tRNA | 149 | 109 | 0.066577 | 0.06745 |
| MT-Inter | 19 | 21 | 0.00849 | 0.012995 |
| Total | 2232 | 1610 | 1 | 1 |

**Separate Excel files**

**Supplemental Table 6**. Annotation of heteroplasmic mutations identified in the Framingham Heart Study (FHS_annotation_heteroplasmy.csv)

**Supplemental Table 7**. Annotation of heteroplasmic mutations identified in the Jackson Heart Study (JHS_annotation_heteroplasmy.csv)

**Supplemental Table 8**. Annotation of heteroplasmic mutations identified in the Cleveland Family Study (CFS_annotation_heteroplasmy.csv)

**Supplemental Table 9. Comparison of heteroplasmic mutations identified using rCRS and RSRS.**

A. **Nine mtDNA heteroplasmic loci were distributed differently in FHS participants.**

| **mtID** | **rCRS** | **RSRS** | **diff** |
| --- | --- | --- | --- |
| mt73 | 28 | 29 | 1 |
| mt12705 | 285 | 280 | -5 |
| mt13105 | 186 | 185 | -1 |
| mt16129 | 179 | 171 | -8 |
| mt16189 | 149 | 323 | 174 |
| mt16223 | 92 | 93 | 1 |
| mt16230 | 24 | 23 | -1 |
| mt16311 | 119 | 118 | -1 |
| mt16519 | 96 | 95 | -1 |

rCRS, the number of participants with a mutation using rCRS; RSRS, the number of participants with a mutation using RSRS.

B. **Fifty-nine mtDNA heteroplasmic loci were distributed differently in JHS participants.**

| mtID | rCRS | RSRS | diff |  | mtID | rCRS | RSRS | diff |
| --- | --- | --- | --- | --- | --- | --- | --- | --- |
| mt152 | 22 | 23 | 1 |  | mt15500 | 2 | 3 | 1 |
| mt195 | 18 | 17 | -1 |  | mt15940 | 29 | 30 | 1 |
| mt464 | 36 | 37 | 1 |  | mt16092 | 10 | 11 | 1 |
| mt515 | 297 | 299 | 2 |  | mt16183 | 619 | 620 | 1 |
| mt523 | 4 | NA | NA |  | mt16187 | 344 | 359 | 15 |
| mt545 | 101 | 100 | -1 |  | mt16189 | 128 | 339 | 211 |
| mt564 | 83 | 82 | -1 |  | mt16192 | 223 | 225 | 2 |
| mt567 | 80 | 79 | -1 |  | mt16218 | 31 | 32 | 1 |
| mt1260 | NA | 1 | NA |  | mt16223 | 112 | 141 | 29 |
| mt1438 | 7 | 6 | -1 |  | mt16230 | 67 | 65 | -2 |
| mt2885 | 7 | 8 | 1 |  | mt16249 | 69 | 71 | 2 |
| mt5237 | 4 | 5 | 1 |  | mt16259 | 4 | 5 | 1 |
| mt5261 | 4 | 5 | 1 |  | mt16263 | 49 | 51 | 2 |
| mt5821 | 3 | 4 | 1 |  | mt16264 | 13 | 14 | 1 |
| mt7256 | 99 | 100 | 1 |  | mt16274 | 25 | 26 | 1 |
| mt7686 | NA | 1 | NA |  | mt16278 | 25 | 26 | 1 |
| mt8154 | NA | 1 | NA |  | mt16284 | 6 | 7 | 1 |
| mt8170 | NA | 1 | NA |  | mt16288 | 7 | 8 | 1 |
| mt8395 | 1 | 2 | 1 |  | mt16290 | 5 | 6 | 1 |
| mt9221 | 12 | 13 | 1 |  | mt16293 | 26 | 27 | 1 |
| mt10308 | 1 | 2 | 1 |  | mt16294 | 39 | 40 | 1 |
| mt10953 | 66 | 67 | 1 |  | mt16301 | 20 | 21 | 1 |
| mt12275 | 1 | 2 | 1 |  | mt16309 | 62 | 64 | 2 |
| mt12812 | NA | 1 | NA |  | mt16311 | 65 | 63 | -2 |
| mt13105 | 61 | 60 | -1 |  | mt16355 | 23 | 24 | 1 |
| mt13781 | 11 | 12 | 1 |  | mt16356 | 22 | 23 | 1 |
| mt13803 | 9 | 10 | 1 |  | mt16368 | 13 | 14 | 1 |
| mt14311 | 3 | 4 | 1 |  | mt16399 | 15 | 16 | 1 |
|  |  |  |  |  | mt16527 | 49 | 50 | 1 |

rCRS, the number of participants with a mutation using rCRS; RSRS, the number of participants with a mutation using RSRS.

**Supplemental Table 10**. Comparison of concordance rate of heteroplasmic mutations identified between rCRS and RSRS

| **Pair type** | **rCRS** | **RSRS** |
| --- | --- | --- |
| FHS | | |
| Mother-offspring | 21.00% | 21.90% |
| Sibling-sibling | 17.70% | 18.20% |
| Extended maternal | 16.40% | 17.60% |
| Father-offspring | 1.50% | 1.80% |
| Unrelated of L haplogroup | 1.70% | 1.70% |
| Unrelated mixed haplogroup | 1.10% | 1.50% |
| JHS | | |
| Mother-offspring | 40.80% | 39.90% |
| Sibling-sibling | 36.00% | 36.20% |
| Extended maternal | 30.20% | 30.90% |
| Father-offspring | 4.50% | 6.10% |
| Unrelated of L haplogroup | 8.70% | 8.70% |
| Unrelated mixed haplogroup | 6.20% | 7.20% |

**Supplemental Table 11**. The association analysis between the burden of heteroplasmic mutations and sex

|  | FHS, beta/SE (p value) | | |
| --- | --- | --- | --- |
| Sample n | 4036 (pooled) | 2555 (with cell counts) | 2555 (with cell counts) |
| p value | -0.035/0.031 (0.26) | -0.026/0.038 (0.5) | 0.019/0.040 (0.63) |
|  | JHS, beta/SE (p value) | | |
| Sample n | 3404 (pooled) | 2737 (with cell counts) | 2737 (with cell counts) |
| p value | 0.12/0.035 (0.0004) | 0.13/0.04 (0.001) | 0.12/0.04 (0.002) |

The heteroplasmy burden residuals were obtained after regressing heteroplasmy counts on age and batch (pooled sample, n=4036 in the FHS, n=3404 in the JHS) and cell counts (n=2555 in the FHS and n=2737 in the JHS).

**Supplemental Table 12**. The association analysis between the burden of heteroplasmic mutations and white blood cells

| Cell counts | **FHS (n=2,555)** | | **JHS(n=2,737)** | |
| --- | --- | --- | --- | --- |
|  | p | R2 | p | R2 |
| WBC | 2.7E-07 | 0.01 | 0.0013 | 0.003 |
| NE | 2.5E-06 | 0.008 | n.a. | n.a. |
| LY | 1.2E-05 | 0.007 | 0.007 | 0.004 |
| EO | 4.0E-02 | 0.001 | 0.6 | 0 |
| BA | 2.5E-03 | 0.003 | 0.3 | 0 |
| Joint | 1.1E-07 | 0.014 | 0.001 | 0.005 |

WBC, white blood cell count; NE, neutrophil; LY, lymphocyte; EO, eosinophil; BA, basophil; joint, white blood cell counts and differentials jointly in the model. Heteroplasmy burden residuals were obtained by regressing heteroplasmy burden on age, sex, and batch effect.

**Supplemental Table 13**. The association analysis between the burden of heteroplasmic mutations and mtDNA haplogroups

A. mtDNA haplogroups in European Americans

| **mtDNA  haplogroup** | **FHS** | | **CFS EA** | |
| --- | --- | --- | --- | --- |
|  | **N** | **%** | **N** | **%** |
| H | 1881 | 46.31 | 269 | 44.98 |
| U | 522 | 12.85 | 70 | 11.71 |
| J | 365 | 8.99 | 53 | 8.86 |
| T | 350 | 8.62 | 78 | 13.04 |
| K | 341 | 8.39 | 47 | 7.86 |
| I | 136 | 3.35 | 7 | 1.17 |
| V | 112 | 2.76 | 42 | 7.02 |
| W | 87 | 2.14 | 12 | 2.00 |
| X | 82 | 2.02 | 9 | 1.51 |
| R | 74 | 1.82 | 2 | 0.33 |
| N | 34 | 0.84 | 5 | 0.84 |
| M | 31 | 0.76 | 0 | 0 |
| L | 30 | 0.74 | 2 | 0.33 |
| Others | 17 | 0.42 | 2 | 0.33 |

B. mtDNA haplogroups in African Americans

| **mtDNA haplogroups** | **JHS** | | **CFS AA** | |
| --- | --- | --- | --- | --- |
|  | **N** | **%** | **N** | **%** |
| L0 | 170 | 4.99 | 23 | 3.65 |
| L1 | 718 | 21.1 | 147 | 23.33 |
| L2 | 1004 | 29.49 | 164 | 26.03 |
| L3 | 1340 | 39.36 | 248 | 39.37 |
| L4 | 42 | 1.23 | 5 | 0.79 |
| Others | 130 | 3.82 | 40 | 6.34 |

C. **The Framingham Heart Study:** Pair-wise comparison of heteroplasmy burden between mtDNA haplogroups

|  | **H** | **I** | **J** | **K** | **M** | **N** | **R** | **T** | **U** | **V** | **W** | **X** | **others** |
| --- | --- | --- | --- | --- | --- | --- | --- | --- | --- | --- | --- | --- | --- |
| **H** |  | -8.59 | -2.00 | -2.63 | -5.33 | -2.92 | -8.86 | -1.88 | -6.09 | 0.92 | 0.08 | -5.63 | -4.77 |
|  |  | <.0001 | 1 | 0.6684 | <.0001 | 0.2721 | <.0001 | 1 | <.0001 | 1 | 1 | <.0001 | 0.0001 |
| **I** | 8.59 |  | 6.47 | 6.02 | -1.16 | 1.28 | -2.00 | 6.49 | 4.80 | 7.57 | 5.65 | 0.88 | 0.30 |
|  | <.0001 |  | <.0001 | <.0001 | 1 | 1 | 1 | <.0001 | 0.0001 | <.0001 | <.0001 | 1 | 1 |
| **J** | 2.00 | -6.47 |  | -0.53 | -4.58 | -2.20 | -7.35 | 0.07 | -2.76 | 2.10 | 1.04 | -4.28 | -3.82 |
|  | 1 | <.0001 |  | 1 | 0.0004 | 1 | <.0001 | 1 | 0.4545 | 1 | 1 | 0.0015 | 0.0104 |
| **K** | 2.63 | -6.02 | 0.53 |  | -4.37 | -1.97 | -7.00 | 0.60 | -2.13 | 2.54 | 1.37 | -3.93 | -3.56 |
|  | 0.6684 | <.0001 | 1 |  | 0.001 | 1 | <.0001 | 1 | 1 | 0.8724 | 1 | 0.0068 | 0.0298 |
| **M** | 5.33 | 1.16 | 4.58 | 4.37 |  | 1.91 | -0.23 | 4.60 | 3.66 | 5.38 | 4.71 | 1.67 | 1.22 |
|  | <.0001 | 1 | 0.0004 | 0.001 |  | 1 | 1 | 3E-04 | 0.0203 | <.0001 | 0.0002 | 1 | 1 |
| **N** | 2.92 | -1.28 | 2.20 | 1.97 | -1.91 |  | -2.56 | 2.22 | 1.18 | 3.10 | 2.56 | -0.60 | -0.86 |
|  | 0.2721 | 1 | 1 | 1 | 1 |  | 0.817 | 1 | 1 | 0.1522 | 0.8158 | 1 | 1 |
| **R** | 8.86 | 2.00 | 7.35 | 7.00 | 0.23 | 2.56 |  | 7.37 | 6.04 | 8.29 | 6.73 | 2.57 | 1.81 |
|  | <.0001 | 1 | <.0001 | <.0001 | 1 | 0.817 |  | <.0001 | <.0001 | <.0001 | <.0001 | 0.8044 | 1 |
| **T** | 1.88 | -6.49 | -0.07 | -0.60 | -4.60 | -2.22 | -7.37 |  | -2.80 | 2.03 | 0.99 | -4.30 | -3.85 |
|  | 1 | <.0001 | 1 | 1 | 0.0003 | 1 | <.0001 |  | 0.3979 | 1 | 1 | 0.0013 | 0.0094 |
| **U** | 6.09 | -4.80 | 2.76 | 2.13 | -3.66 | -1.18 | -6.04 | 2.80 |  | 4.54 | 2.70 | -2.83 | -2.67 |
|  | <.0001 | 0.0001 | 0.4545 | 1 | 0.0203 | 1 | <.0001 | 0.398 |  | 0.0004 | 0.5409 | 0.3682 | 0.5941 |
| **V** | -0.92 | -7.57 | -2.10 | -2.54 | -5.38 | -3.10 | -8.29 | -2.03 | -4.54 |  | -0.45 | -5.41 | -4.79 |
|  | 1 | <.0001 | 1 | 0.8724 | <.0001 | 0.1522 | <.0001 | 1 | 0.0004 |  | 1 | <.0001 | 0.0001 |
| **W** | -0.08 | -5.65 | -1.04 | -1.37 | -4.71 | -2.56 | -6.73 | -0.99 | -2.70 | 0.45 |  | -4.21 | -3.97 |
|  | 1 | <.0001 | 1 | 1 | 0.0002 | 0.8158 | <.0001 | 1 | 0.5409 | 1 |  | 0.002 | 0.0058 |
| **X** | 5.63 | -0.88 | 4.28 | 3.93 | -1.67 | 0.60 | -2.57 | 4.30 | 2.83 | 5.41 | 4.21 |  | -0.39 |
|  | <.0001 | 1 | 0.0015 | 0.0068 | 1 | 1 | 0.804 | 0.001 | 0.3682 | <.0001 | 0.002 |  | 1 |
| **Others** | 4.77 | -0.30 | 3.82 | 3.56 | -1.22 | 0.86 | -1.81 | 3.85 | 2.67 | 4.79 | 3.97 | 0.39 |  |
|  | 0.0001 | 1 | 0.0104 | 0.0298 | 1 | 1 | 1 | 0.009 | 0.5941 | 0.0001 | 0.0058 | 1 |  |

D. **The Jackson Heart Study:** Pair-wise comparison of heteroplasmy burden between mtDNA haplogroups

|  | **L0** | **L1** | **L2** | **L3** | **L4** | **Others** |
| --- | --- | --- | --- | --- | --- | --- |
| **L0** |  | -1.10 | 7.97 | 6.17 | 0.19 | 7.69 |
|  |  | 1 | <.0001 | <.0001 | 1 | <.0001 |
| **L1** | 1.10 |  | 15.44 | 12.94 | 0.80 | 9.66 |
|  | 1 |  | <.0001 | <.0001 | 1 | <.0001 |
| **L2** | -7.97 | -15.44 |  | -3.86 | -3.99 | 3.27 |
|  | <.0001 | <.0001 |  | 0.0017 | 0.001 | 0.0163 |
| **L3** | -6.17 | -12.94 | 3.86 |  | -2.99 | 4.70 |
|  | <.0001 | <.0001 | 0.0017 |  | 0.0421 | <.0001 |
| **L4** | -0.19 | -0.80 | 3.99 | 2.99 |  | 5.29 |
|  | 1 | 1 | 0.001 | 0.0421 |  | <.0001 |
| **Others** | -7.69 | -9.66 | -3.27 | -4.70 | -5.29 |  |
|  | <.0001 | <.0001 | 0.0163 | <.0001 | <.0001 |  |

E. The least square means of the heteroplasmy burden in mtDNA haplogroups

| **FHS** | | | |
| --- | --- | --- | --- |
| **mtDNA haplogroup** | **Least square mean** | **SE** | **Count (%)** |
| H | -0.14 | 0.02 | 1747 ( 43.82 ) |
| U | 0.16 | 0.04 | 514 ( 12.89 ) |
| J | -0.03 | 0.05 | 359 ( 9 ) |
| T | -0.03 | 0.05 | 346 ( 8.68 ) |
| K | 0.01 | 0.05 | 337 ( 8.45 ) |
| V | -0.21 | 0.07 | 209 ( 5.24 ) |
| I | 0.61 | 0.08 | 133 ( 3.34 ) |
| W | -0.15 | 0.10 | 86 ( 2.16 ) |
| X | 0.49 | 0.11 | 79 ( 1.98 ) |
| R | 0.90 | 0.11 | 72 ( 1.81 ) |
| M | 0.85 | 0.18 | 28 ( 0.7 ) |
| N | 0.37 | 0.17 | 32 ( 0.8 ) |
| others | 0.56 | 0.15 | 45 ( 1.13 ) |
| **JHS** | | | |
| **mtDNA haplogroup** | **Least square mean** | **SE** | **count (%)** |
| L0 | 0.39 | 0.07 | 170 ( 4.99 ) |
| L1 | 0.48 | 0.04 | 718 ( 21.09 ) |
| L2 | -0.25 | 0.03 | 1004 ( 29.49 ) |
| L3 | -0.09 | 0.03 | 1389 ( 40.8 ) |
| L4 | 0.36 | 0.15 | 42 ( 1.23 ) |
| Others | -0.61 | 0.11 | 81 ( 2.38 ) |

**Supplemental Figure 14. Meta-analysis of EA and AA participants for genome-wide association analysis of mtDNA heteroplasmic mutation burden.**

Separate Excel

**Supplemental Table 15. Previous GWAS loci that are significantly related to heteroplasmic burden**

| **SNPS** | **Chr:pos** | **MAPPED_GENE** | **CONTEXT** | **risk_AF** | **Phenotype** | **P.VALUE** | **Total N** | **P.VALUE** | **PUBMEDID** | **JOURNAL** |
| --- | --- | --- | --- | --- | --- | --- | --- | --- | --- | --- |
| rs11826064 | 11:50818695 | AC024405.1 –  OR4C50P | Intergenic | NR | Depressed affect | 7.00E-10 | 358 K EA |  | 29942085 | Nat Genet |
| rs1814175* | 11:49537620 | AC135977.1 –  AC136759.1 | Intergenic | 0.34 | Height | 2.00E-08 | 184 K EA |  | 20881960 | Nature |
| rs691329 | 11:50428834 | AC024405.1 –  OR4C50P | Intergenic | NR | Creatinine levels | 2.00E-08 | 142 K Jap |  | 29403010 | Nat Genet |
| rs691329 | 11:50428834 | AC024405.1 –  OR4C50P | Intergenic | NR | Glomerular filtration rate | 3.00E-08 | 144 K Jap |  | 29403010 | Nat Genet |
| rs2727020 | 11:49089855 | UBTFL7 –  AC084851.2 | Intergenic | 0.68 | Coronary artery disease | 8.00E-06 | 88.7k/424.5k  case/control |  | 29212778 | Circ Res |
| rs7482876* | 11:49739934 | AC136759.1 | Intron | 0.33 | Feeling fed-up | 4.00E-08 | 375 K EA |  | 29500382 | Nat Commun |
| rs4309155* | 11:50106798 | GTF2IP11 –  AC109635.5 | Intergenic | 0.34 | Feeling fed-up | 4.00E-08 | 375 K EA |  | 29500382 | Nat Commun |
| rs10794221* | 11:50265244 | AC109635.1 –  AC109635.6 | Intergenic | 0.35 | Feeling fed-up | 8.00E-09 | 375 K EA |  | 29500382 | Nat Commun |
| rs692360 | 11:50464323 | AC024405.1 –  OR4C50P | Intergenic | 0.35 | Feeling fed-up | 2.00E-09 | 375 K EA |  | 29500382 | Nat Commun |
| rs11532157 | 11:50813515 | AC024405.1 –  OR4C50P | Intergenic | 0.30 | Feeling fed-up | 5.00E-10 | 375 K EA |  | 29500382 | Nat Commun |
| rs3862341 | 11:49275575 | FOLH1 –  AC118942.1 | Intergenic | NR | Hair color | 3.00E-07 | 375 K EA |  | 30595370 | Am J Hum Genet |
| rs4980470 | 11:50665628 | AC024405.1 –  OR4C50P | Intergenic | 0.59 | Systolic blood pressure | 1.00E-08 | 776 K EA+multi |  | 30578418 | Nat Genet |
| rs10769601* | 11:49660329 | AC136759.1 | Intron | NR | Height | 6.00E-12 | 458 K EA |  | 30595370 | Am J Hum Genet |

*, These SNPs are significant in meta-analysis of FHS and JHS GWAS of heteroplasmic burden

**Supplemental Table 16. The gene expression of protein tyrosine phosphatase receptor type J is long cis-eQTLs for GWAS loci**

Separate Excel file

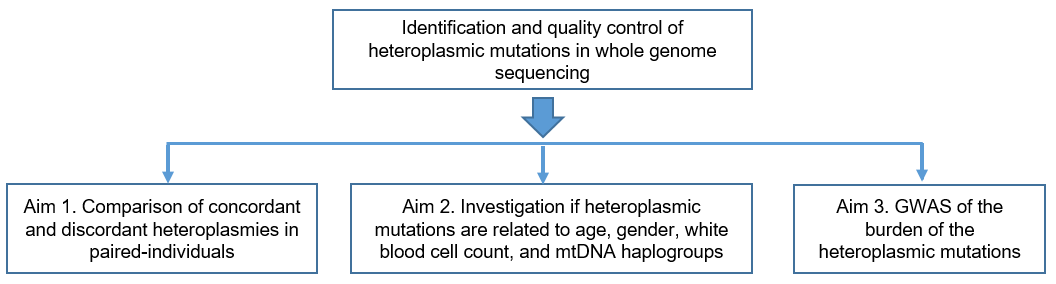

**Supplemental Figure 1**. Study design flow-chart and main aims in this study

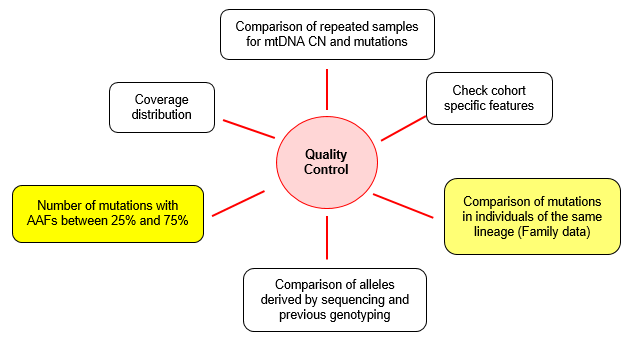

**Supplemental Figure 2**. Comprehensive quality control procedures for heteroplasmic mutations

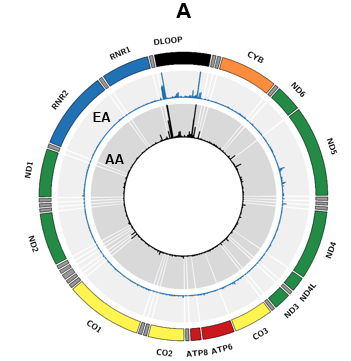

**Supplemental Figure 3**. Description of mtDNA heteroplasmic mutations in the Cleveland Family Study. EA, participants of European descent; AA, participants of African descent.

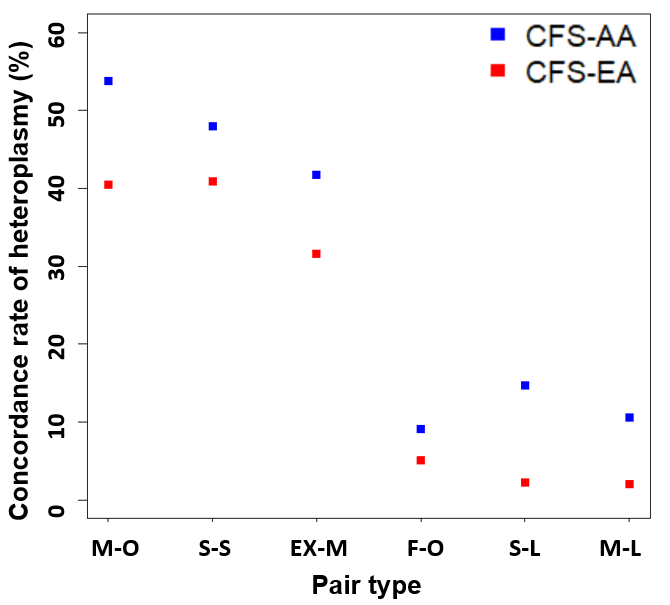

**Supplemental Figure 4**. **Concordance rate of heteroplasmic mutations between paired individuals in CFS AA and EA participants.** M-O, mother-offspring; S-S, sibling-sibling; EX-M, distantly related maternal pairs; F-O, father-offspring; S-L, unrelated pairs in the same mtDNA haplogroup; M-L, unrelated pairs of mixed mtDNA haplogroups.

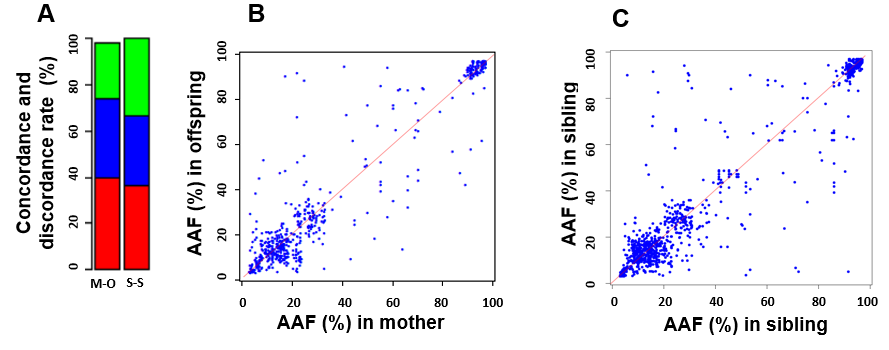

**Supplemental Figure 5. Concordant heteroplasmic mutations in paired JHS individuals**. **A**. Concordant (red color) and discordant (green and blue colors) heteroplasmic mutations between mother-offspring (M-O) and sibling-sibling (S-S) pairs, in mother-offspring bar, the blue portion is the proportion of discordant heteroplasmic mutations in mothers only and the green portion is the proportion of discordant mutations in offspring only, the green and blue proportions in the sibling-sibling bar represent the proportion of mutations in sibling pairs; **B**. Comparison of alternative allele fractions between mother and offspring pairs; **C**. Comparison of alternative allele between sibling-sibling pairs.

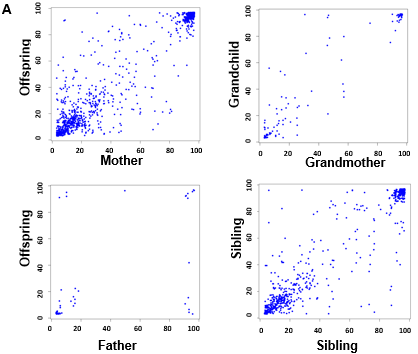

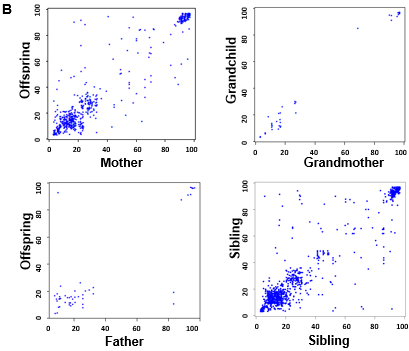

**Supplemental Figure 6**. Comparison of alternative allele fractions between paired individuals in (**A**) the Framingham Heart Study, and (**B**) the Jackson Heart Study. We compared PAA in four types of pairs: mother-offspring, grandmother-grandchild, father-offspring, and sibling-sibling pairs.

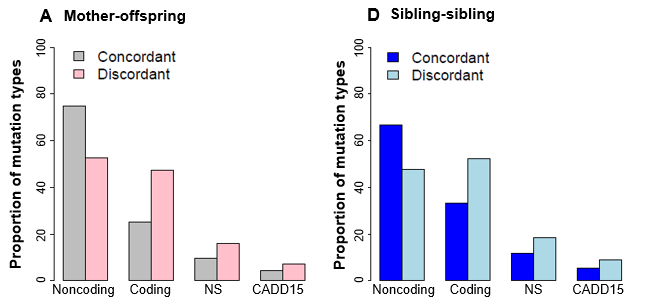

**Supplemental Figure 7**. **Discordant heteroplasmic mutations are more likely to be deleterious than concordant heteroplasmic mutations**. The proportion of concordant heteroplasmic mutations in noncoding, coding regions, being nonsynonymous (NS), and being deleterious with CADD ≥15 (CADD15) in Mother-offspring and sibling-sibling pairs in the Jackson Heart Study (JHS).

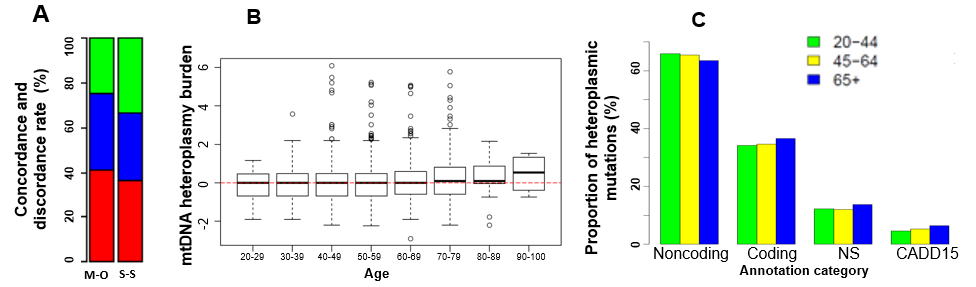

**Supplemental Figure 8**. **Heteroplasmy and age in participants of African American**. **A**. Concordant (red color) and discordant (green and blue colors) heteroplasmic mutations between mother-offspring (M-O) and sibling-sibling (S-S) pairs, in mother-offspring bar, the blue portion is the proportion of discordant heteroplasmic mutations in mothers only and the green portion is the proportion of discordant mutations in offspring only, the green and blue proportions in the sibling-sibling bar represent the proportion of mutations in sibling pairs; **B**. The overall burden of mtDNA heteroplasmic mutations in different age groups; **C**. The proportion of heteroplasmic mutations according to functional annotations in individuals of 20-44, 45-64 and 65 or above; Noncoding, the proportion of heteroplasmies in noncoding regions. Coding, the proportion of heteroplasmies in 13 protein-coding regions; NS, the proportion of nonsynonymous in coding heteroplasmies; CADD15, the proportion of possibly deleterious nonsynonymous mutations with CADD≥15 in coding heteroplasmies.

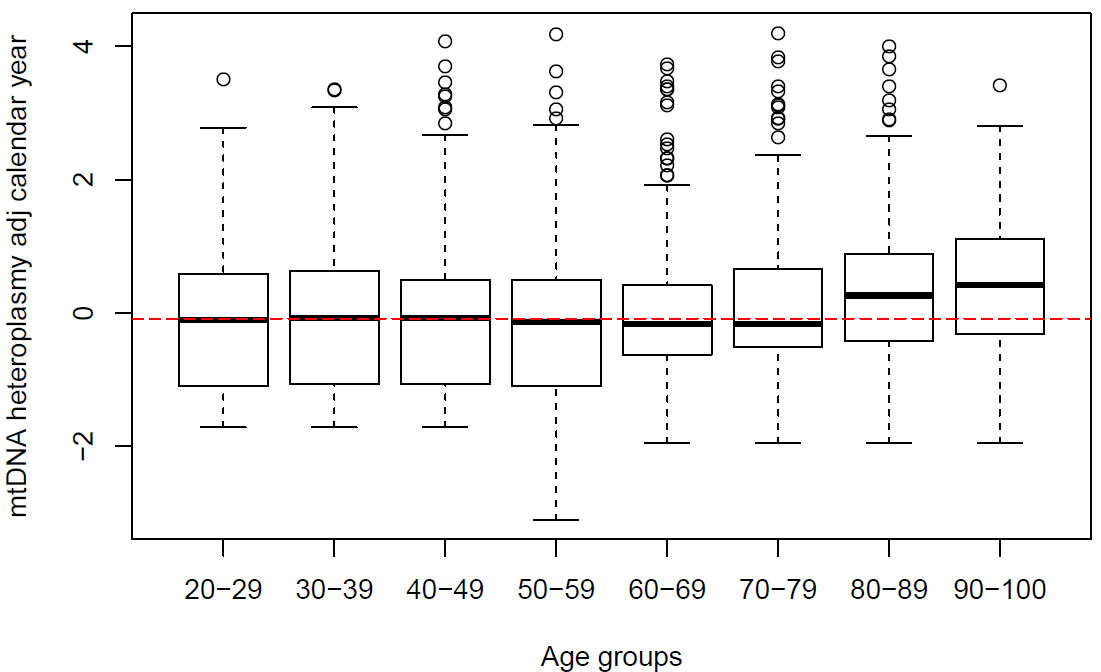

| Agegroup | 20-29 | 30-39 | 40-49 | 50-59 | 60-69 | 70-79 | 80-89 | 90-100 |
| --- | --- | --- | --- | --- | --- | --- | --- | --- |
| N | 110 | 322 | 584 | 815 | 979 | 815 | 448 | 51 |

A. Framingham Heart Study. Heteroplasmy burden residuals were obtained by regressing heteroplasmy burden on batch effect (n=4,036).

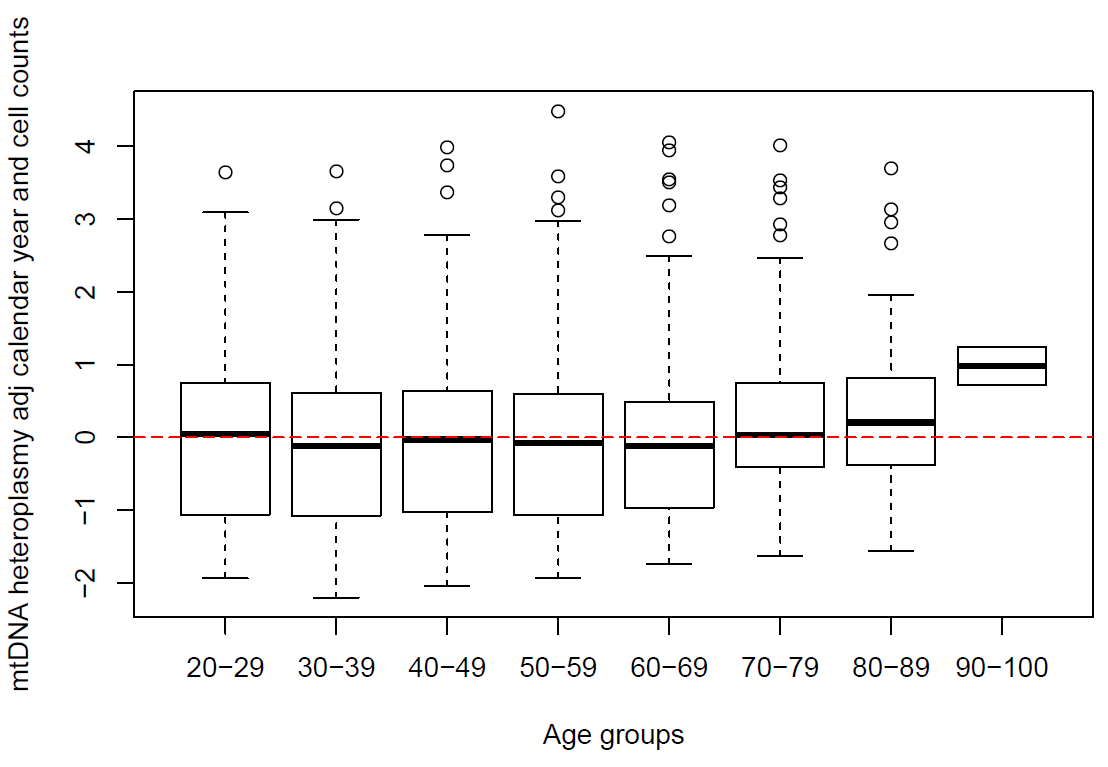

B. Framingham Heart Study. Heteroplasmy burden residuals were obtained by regressing heteroplasmy burden on batch effect and cell counts (n=2,555)

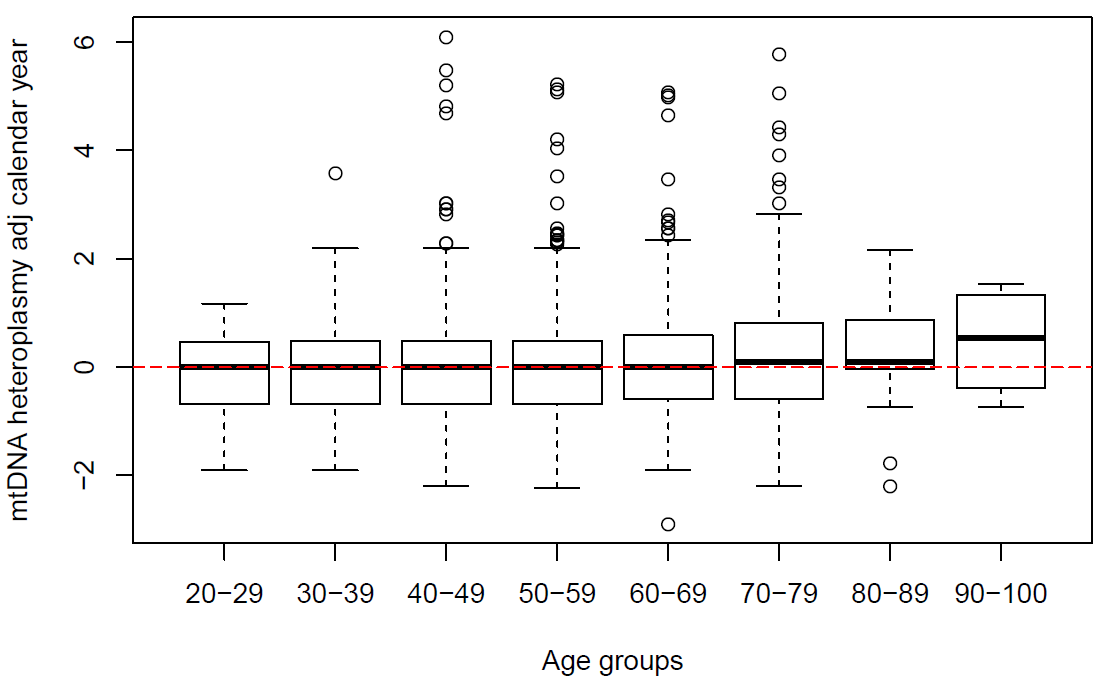

| Agegroup | 20-29 | 30-39 | 40-49 | 50-59 | 60-69 | 70-79 | 80-89 | 90-100 |
| --- | --- | --- | --- | --- | --- | --- | --- | --- |
| N | 75 | 254 | 764 | 810 | 941 | 502 | 54 | 4 |

C. Jackson Heart Study. Heteroplasmy burden residuals were obtained by regressing heteroplasmy burden on batch effect (n=3,404).

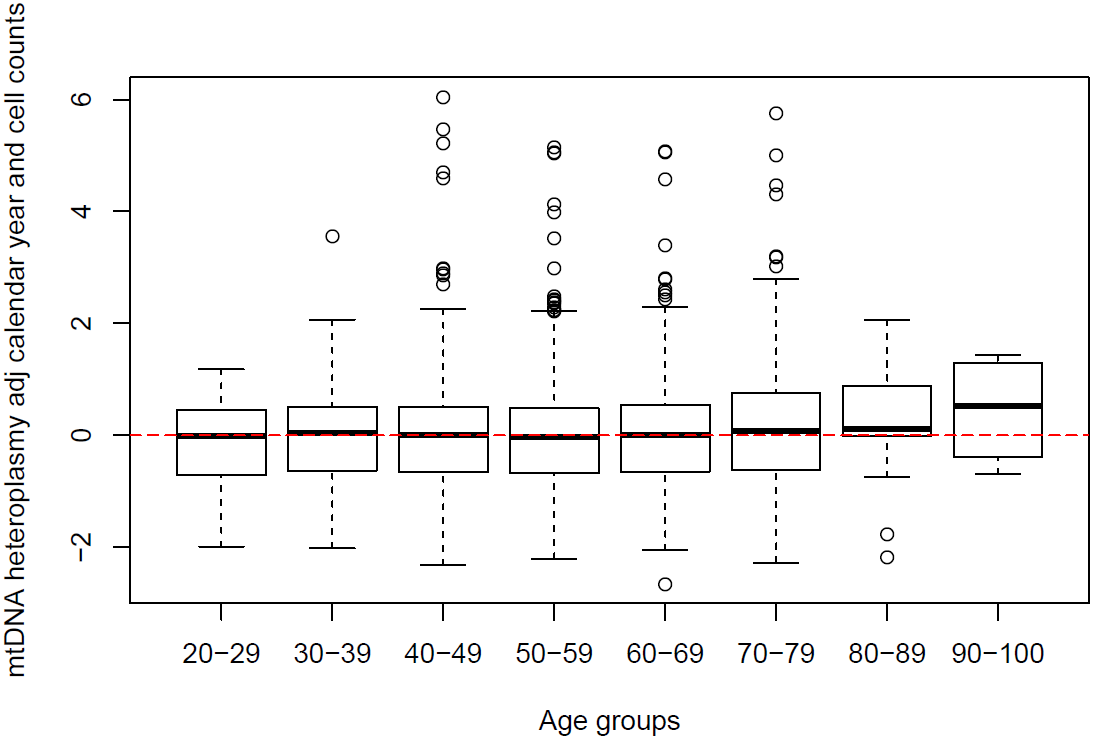

D. Jackson Heart Study. Heteroplasmy burden residuals were obtained by regressing heteroplasmy burden on batch effect and cell counts (n=2,737).

**Supplemental Figure 9**. Heteroplasmy burden and age in the Framingham Heart Study and Jackson Heart Study

**A.**

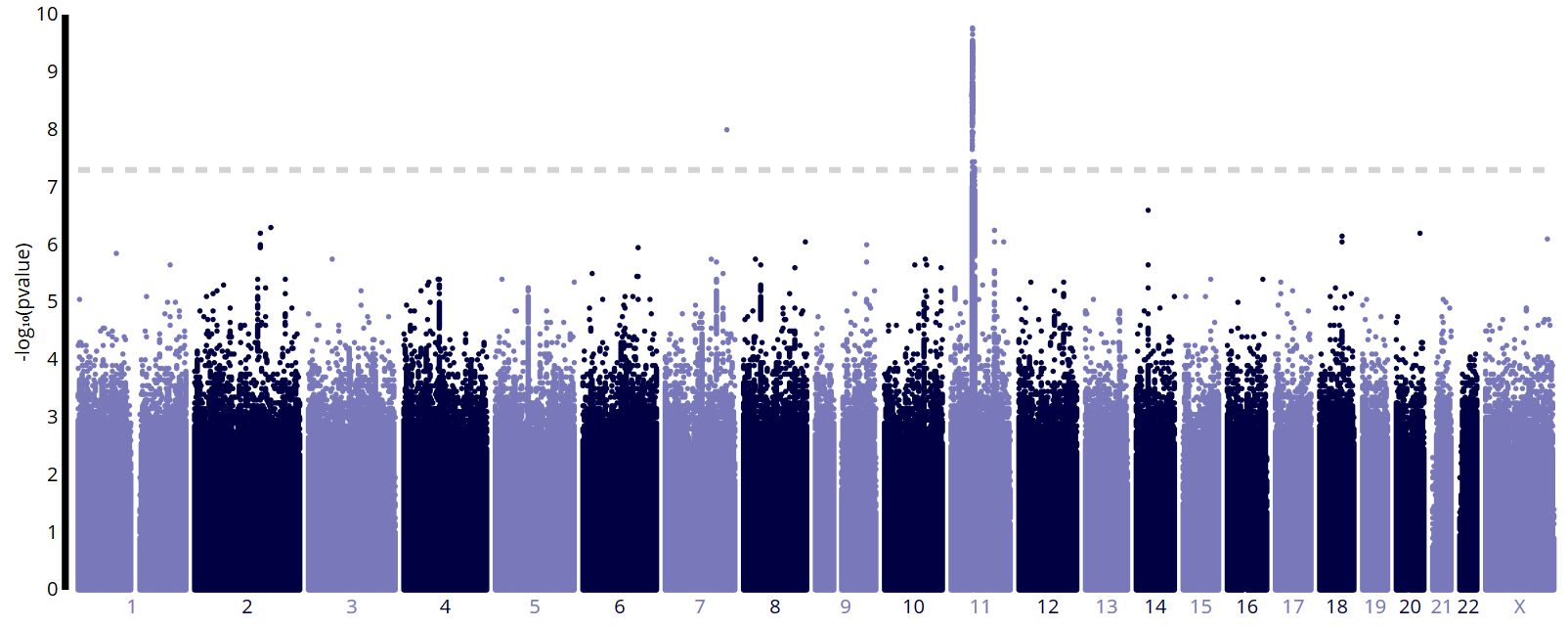

**B**.

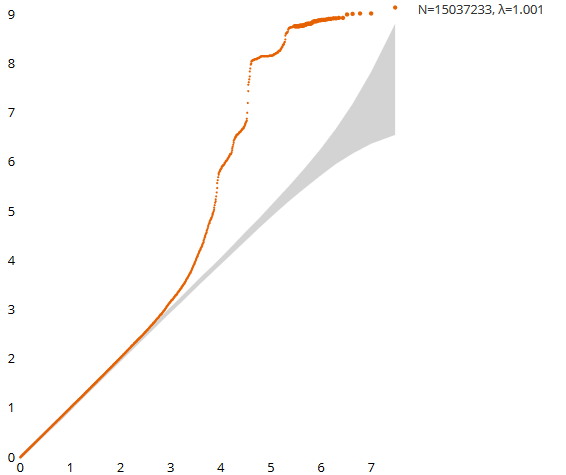

**Supplemental Figure 10**. Genome-wide association analysis of heteroplasmic mutation burden in the mtDNA in the Framingham Heart Study participants. A. Manhattan plot. B. Q-Q plot.

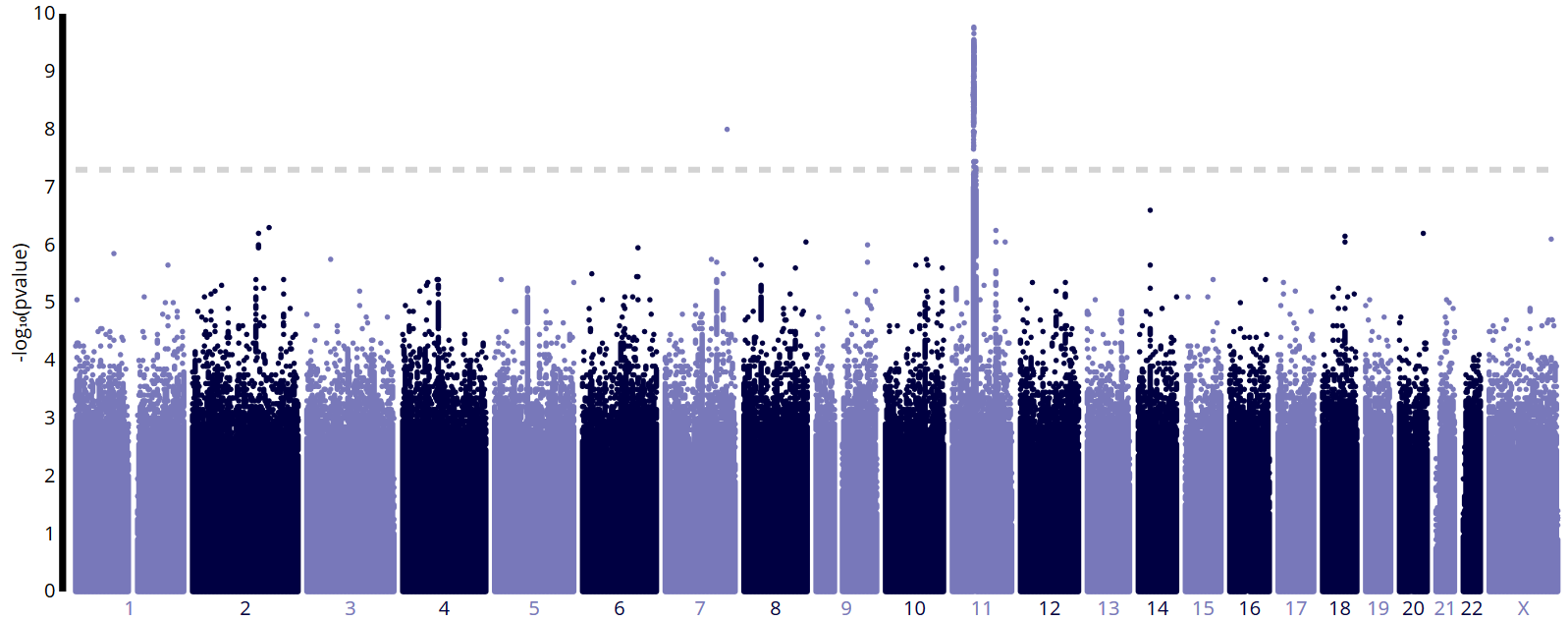

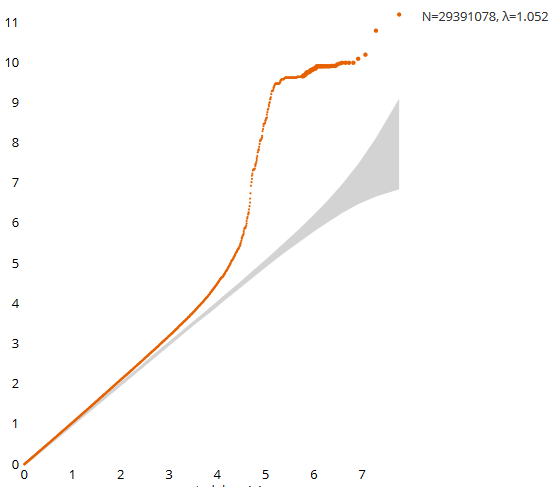

**Supplemental Figure 11**. Genome-wide association analysis of heteroplasmic mutation burden in the mtDNA in the Jackson Heart Study participants. A. Manhattan plot. B. Q-Q plot.

**A**

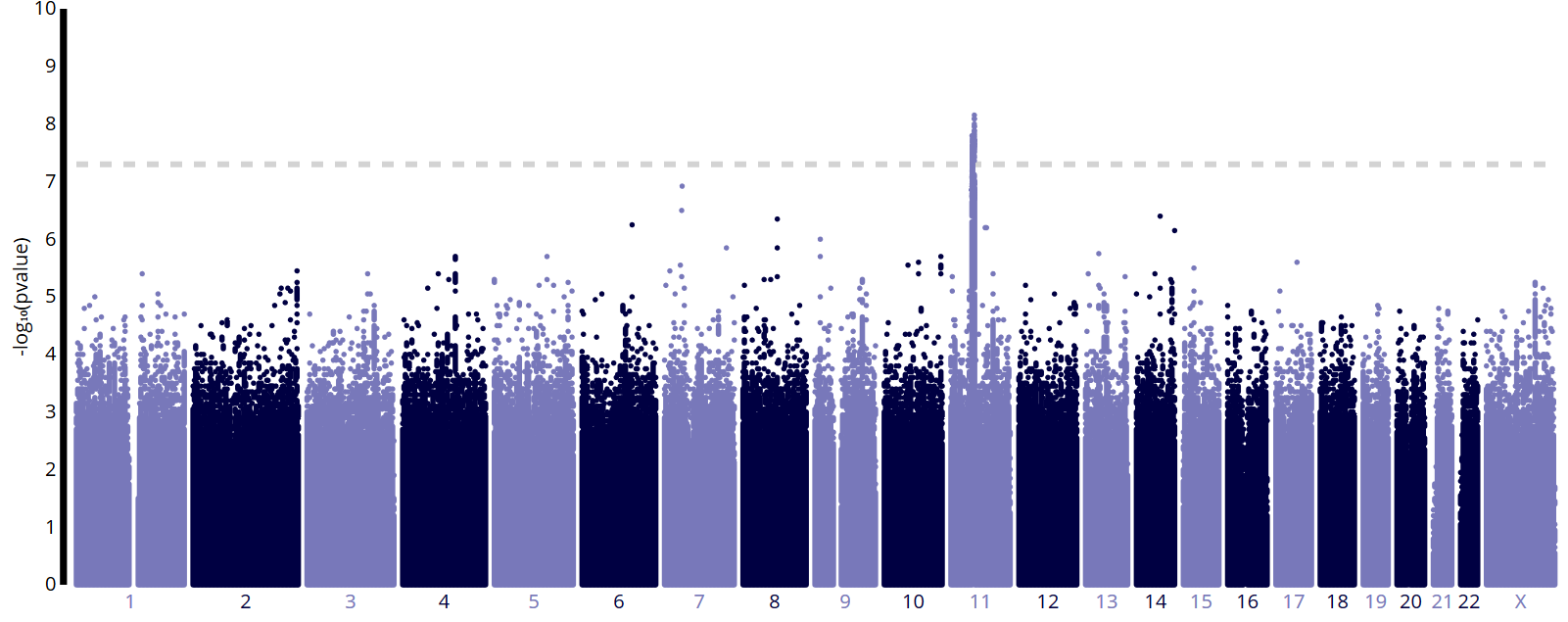

**B.**

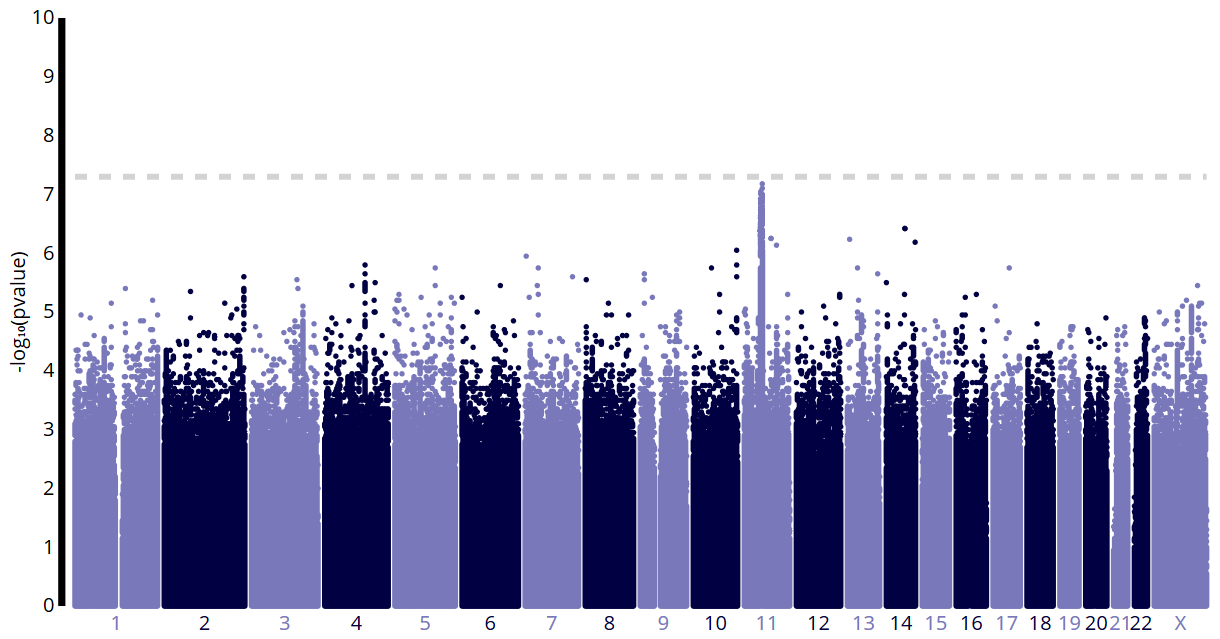

**Supplemental Figure 12**. Genome-wide association analysis of heteroplasmic mutation burden in the same individuals with cell count measurements in the Framingham Heart Study. A. Without cell counts in the model. B. With cell counts in the model.

**A.**
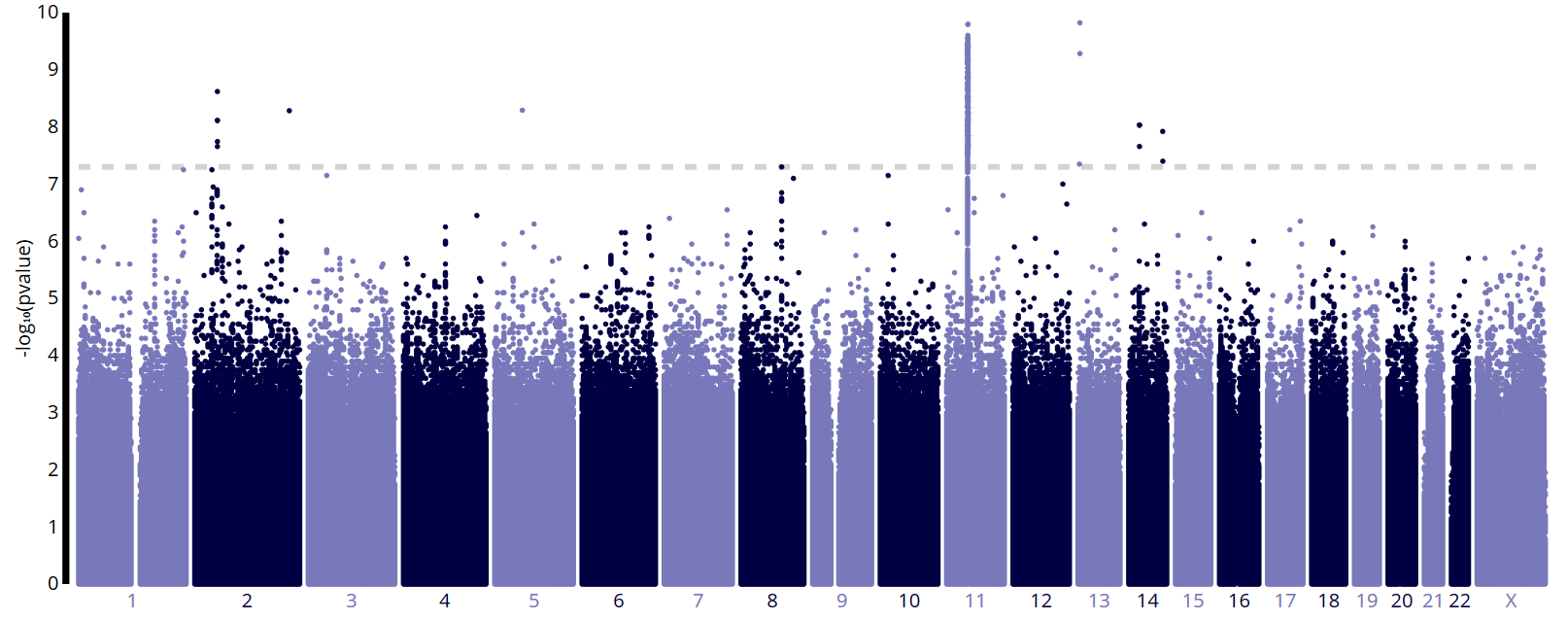

**B.**

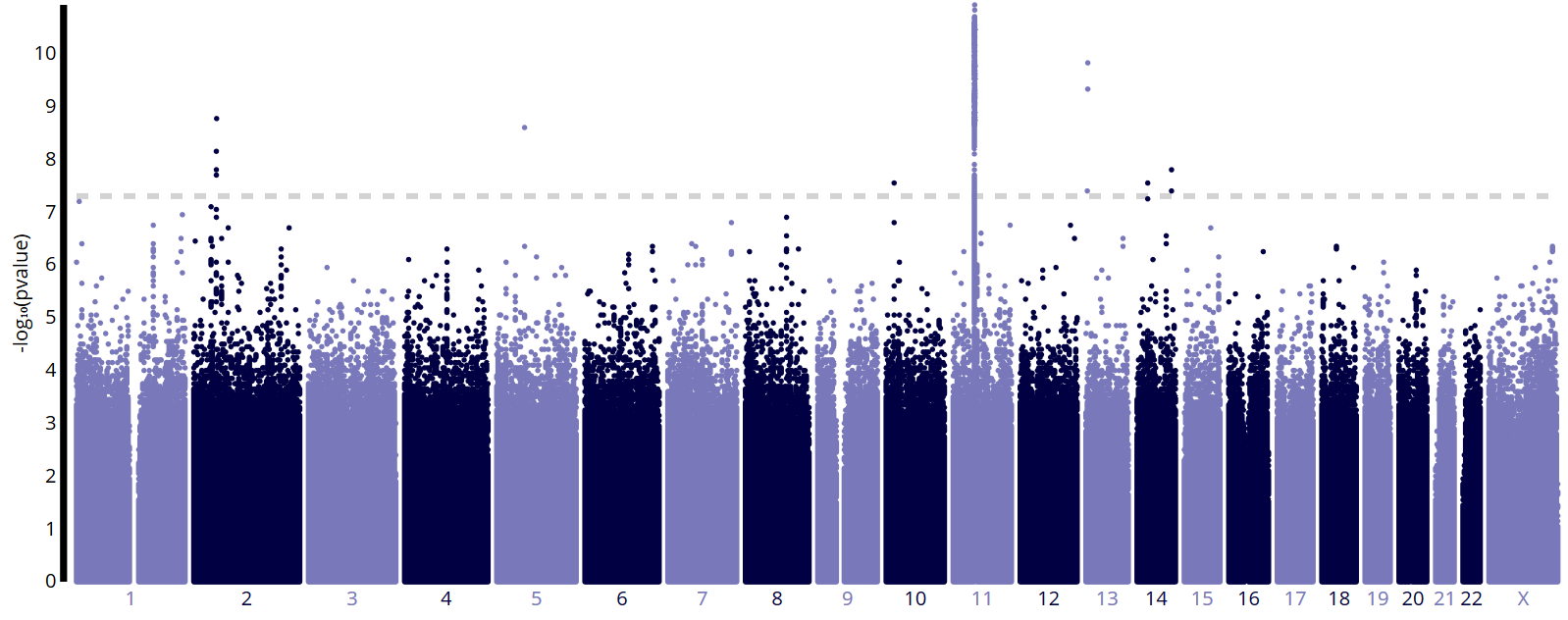

**Supplemental Figure 13**. Genome-wide association analysis of heteroplasmic mutation burden in the same individuals with cell count measurements in the Jackson Heart Study. A. Without cell counts in the model. B. With cell counts in the model.
